# Telmisartan and Lisinopril Show Potential Benefits in Rescuing Cognitive-Behavioral Function Despite Limited Improvements in Neuropathological Outcomes in Tg-SwDI Mice

**DOI:** 10.1101/2025.04.23.650226

**Authors:** Natalia Motzko Noto, Yazmin M. Restrepo, Ariana Hernandez, Chana Vogel, Victoria Pulido-Correa, Shay Moen, Julianna Bonetti, Vedika Chiduruppa, Robert C. Speth, Lisa S. Robison

## Abstract

Cerebral amyloid angiopathy (CAA) is a cerebrovascular disease that results from beta-amyloid (Aβ) accumulation in the vessel walls that is associated with cognitive impairment and other neurological pathologies. There are currently no medications approved to treat CAA. This study investigated whether renin-angiotensin system (RAS)-targeting drugs, commonly prescribed to treat hypertension, can be repurposed to treat CAA, and whether their effects differ by sex. Male and female Tg-SwDI mice were treated for 5 months with sub-depressor doses of either telmisartan [angiotensin II receptor blocker (ARB)] or lisinopril [angiotensin-converting enzyme (ACE) inhibitor] starting at 3 months of age. Blood pressure monitoring was performed 2 and 4 months after the start of treatment, followed by behavior testing at 7 months of age. Histochemical analyses were conducted to determine vasculopathy, Aβ pathology, and neuroinflammation (microgliosis and astrogliosis). Outcomes in drug-treated and untreated Tg-SwDI mice were compared to each other and with wild-type (C57BL/6J) controls. Overall, both drugs were able to rescue some cognitive-behavioral functions; however, no reductions in Aβ levels were observed, and only limited improvements in vascular density and neuroinflammatory markers were detected. Notably, some treatment effects varied with sex, the specific behavioral task, and the brain region analyzed. These findings support the hypothesis that RAS-targeting drugs exert neuroprotective effects through mechanisms beyond blood pressure control offering a promising therapeutic avenue for CAA.

## 1. Introduction

Cerebral amyloid angiopathy (CAA) is a common cerebrovascular disease (DeSimone et al., 2017) in which amyloid-beta (Aβ) peptide aggregates accumulate in the tunica media layer of the walls of cerebral arteries, arterioles, and capillaries of the leptomeninges and cortex; usually surrounding smooth muscle cells (Biffi & Greenberg, 2011; Chwalisz, 2021; DeSimone et al., 2017; Mendel et al., 2013; Weller et al., 2008). As the disease progresses, these smooth muscle cells deteriorate and are replaced by Aβ aggregates (Biffi & Greenberg, 2011; Mendel et al., 2013) with subsequent impairment of cerebral blood flow and blood-brain barrier integrity (Gireud-Goss et al., 2021). CAA is associated with cerebral hemorrhage, and infarcts, leading to neurological deficits (Chwalisz, 2021; Kozberg et al., 2021), including cognitive impairment and dementia (Chwalisz, 2021).

Currently, there are no pharmaceutical agents available that can cure or halt the progression of CAA (Cozza et al., 2023; Saito et al., 2021). Notably, CAA co-exists with Alzheimer’s disease (AD) in ∼90% of AD cases, exacerbating AD pathology and symptoms (Chwalisz, 2021; Saito et al., 2021). Importantly, the presence of significant CAA pathology may preclude AD patients from receiving treatment with anti-amyloid therapies such as monoclonal antibodies due to an increased risk of amyloid-related imaging abnormality (ARIA), including brain swelling (ARIA-E) and/or bleeding (ARIA-H) (Agarwal et al., 2023; Sin et al., 2023; Sveikata et al., 2022). These challenges underscore the urgent need for alternative treatment strategies for CAA, whether it occurs as a singular pathology or in cases in which AD presents with significant CAA.

The Renin-Angiotensin System (RAS) plays a vital role in controlling blood pressure and electrolyte balance (Santos et al., 2018; Speth, 2022). Through its activity in the brain, the RAS can affect cognition, anxiety, depression, and emotional stress (Gebre et al., 2018; Kehoe et al., 2009; Loera-Valencia et al., 2021). In both the vasculature and the brain, the RAS is divided into a classical arm [angiotensin-converting enzyme (ACE)/angiotensin (Ang) II/Ang II receptor type 1 (AT_1_R)] and a counterregulatory arm [ACE-2/Ang-(1-7)/Mas receptor (MasR)]. Effects caused by the activation of each arm in the vasculature and in the brain can be seen in **Figure 1**. Of note, the AT_2_ receptor type which also responds to Ang II, also functions in a counterregulatory manner to the effects mediated by Ang II at the AT_1_R (de Gasparo et al., 2000; Restrepo et al., 2022).

**Figure 1.**
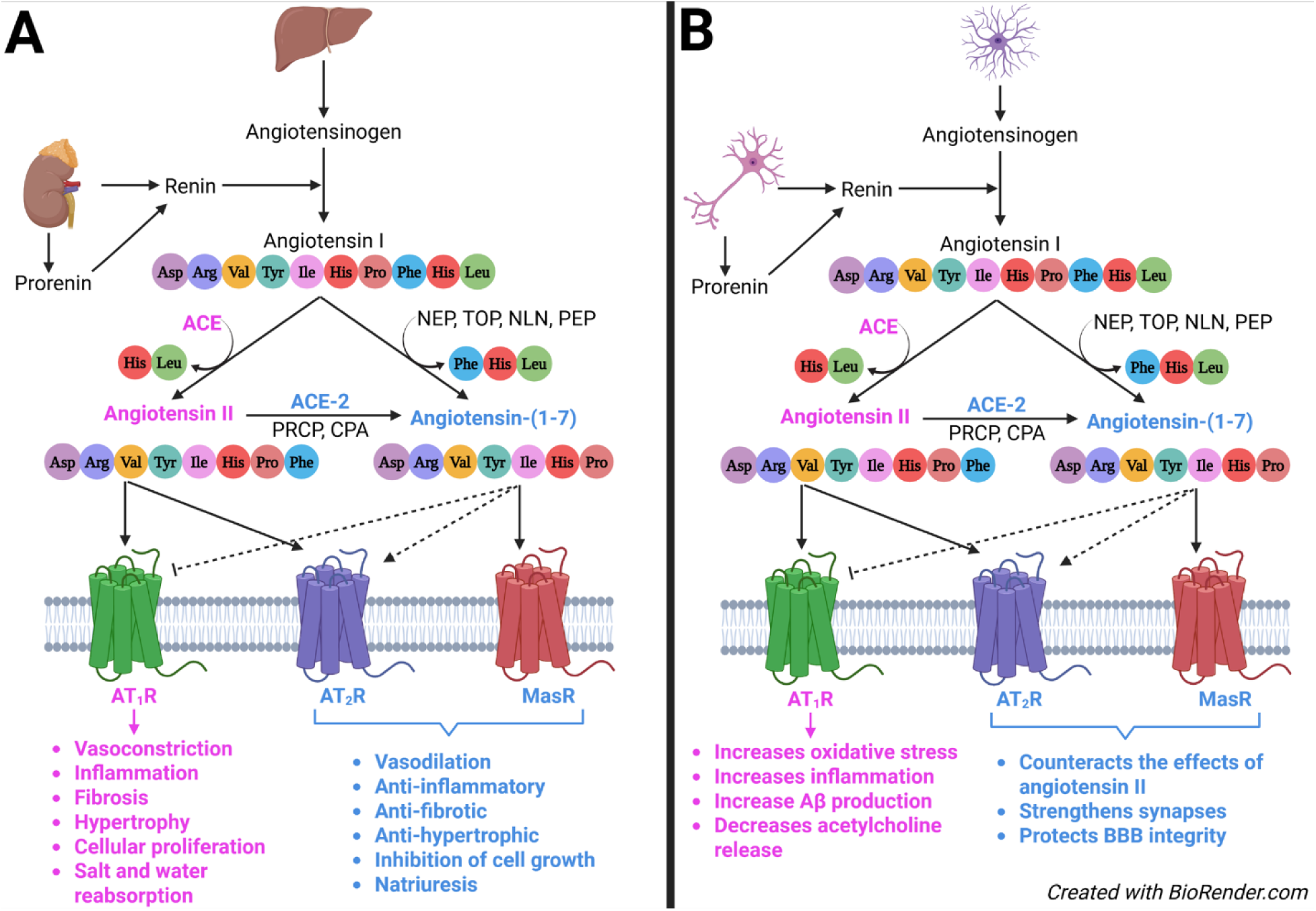
Simplified representation of the activity of the vasculature and the brain RAS. **(A)** In the vasculature, the enzyme renin, as well as its precursor prorenin, are released by the kidneys. Renin cleaves angiotensinogen, giving rise to angiotensin (Ang) I, which is then cleaved by the angiotensin-converting enzyme (ACE), generating Ang II. Ang II activates both Ang II type 1 receptors (AT_1_Rs) and Ang II type 2 receptors (AT_2_Rs), and can be further cleaved by ACE-2, prolylcarboxypeptidase (PRCP), or carboxypeptidase A (CPA) to generate Ang-(1-7), which activates MasRs and AT_2_Rs (to a lesser extent), and inhibits AT_1_Rs, counteracting the effects of Ang II. Alternatively, Ang-(1-7) can also be formed through cleavage of Ang I by neprilysin (NEP), thimet oligopeptidase (TOP), neurolysin (NLN), and prolylendopeptidase (PEP). **(B)** In the brain, angiotensinogen is produced by astroglia (Stornetta et al., 1988) and can be cleaved by renin, prorenin, or renin-like enzymes present in neurons, giving rise to Ang I (Cooper et al., 2022; Husain et al., 1984). Ang I is then cleaved by ACEs, producing Ang II (Strittmatter et al., 1985). Similar to the vasculature RAS, Ang-(1-7) counteracts the effects of Ang II in the brain. *Created in BioRender. Speth, R. (2025)* https://BioRender.com/rg5y89w

The RAS has been identified as a potential target to treat dementia, particularly AD, as brain RAS dysregulation has been observed in both patients and animal models of these diseases (Arregui et al., 1982; Chen et al., 2023; Cosarderelioglu et al., 2022; Miners et al., 2008; Savaskan et al., 2001). Substantial evidence supporting this approach comes from epidemiological studies showing that individuals taking RAS-targeting drugs, specifically those capable of penetrating the blood-brain barrier (BBB), have a reduced risk of all-cause dementia, AD, and vascular contributions to cognitive impairment and dementia, even after controlling for the drugs’ cardiovascular benefits (Dahlof et al., 2002; Rea et al., 2024; Tsukuda et al., 2009; Yasar et al., 2013). Researchers have explored repurposing RAS-targeting drugs, such as angiotensin receptor blockers (ARBs) and ACE inhibitors, to prevent/treat neurodegenerative diseases, particularly AD (Kehoe & Wilcock, 2007; Lee et al., 2023; Ouk et al., 2021). While these studies have shown promising results (Cosarderelioglu et al., 2020; Ribeiro et al., 2020), studies that examine the use of RAS-targeting drugs to address CAA specifically are still lacking.

Due to evidence of RAS dysregulation in neurodegenerative diseases (Arregui et al., 1982; Chen et al., 2023; Cosarderelioglu et al., 2022; Miners et al., 2008; Savaskan et al., 2001) and the demonstrated neuroprotective effects of RAS-targeting drugs (Cosarderelioglu et al., 2020; Ribeiro et al., 2020), we investigated whether RAS-inhibiting drugs (ARBs and ACE inhibitors) are capable of preventing the cognitive decline and neuropathology associated with CAA in Tg-SwDI mice, a mouse model of CAA in early-stage disease; while also determining if the drug effects are sex-specific. We hypothesized that ARBs and ACE inhibitors would have therapeutic value for preserving cognitive-behavioral functions by reducing vascular Aβ protein accumulation and associated neuropathologies.

## 2. Materials and Methods

### 2.1. Animals

Tg-SwDI mice (JAX MMRRC Stock #034843), which exhibit behavioral deficits, amyloid plaque and fibrillar vascular Aβ pathology, and perivascular inflammation (Davis et al., 2004; Robison et al., 2019), were used as a transgenic animal model exhibiting CAA. At approximately 3 months of age, Aβ starts to accumulate primarily around the cerebral vasculature of these mice, with increasing severity over their lifespan (Paul et al., 2018; Rosas-Hernandez et al., 2020; Xu et al., 2014).

Tg-SwDI mice were divided into male and female groups and were age- and sex-matched with C57BL/6J mice, their background strain, which served as wild-type (WT) controls. Males and females were kept in separate cages, with 3-to-5 littermates per cage. Mice were maintained on a reverse 12-hour light cycle (lights on at 8/9 PM, lights off at 8/9AM, varying with daylight-savings time). Temperature and humidity were 68-72°F and 40-60%, respectively. Food and water were available ad libitum.

The Nova Southeastern University Institutional Animal Care and Use Committee approved the experimental procedures used in this study (approval no. 2022.04.LR2). All animal housing and experiments were conducted in strict accordance with the Institutional Guidelines for Care and Use of Laboratory Animals at Nova Southeastern University.

### 2.2. Drug treatment

Tg-SwDI mice were assigned to drug groups to receive sub-depressor doses of either telmisartan [1 mg/kg/day (Washida et al., 2010)] or lisinopril [15 mg/kg/day (Gonzalez-Villalobos et al., 2009; Singh et al., 2013)], dissolved in reverse osmosis water. The concentrations for the drug solutions were calculated every week for each cage based on the average body weight of the mice and the average volume of fluid consumption in the previous week. Treatment lasted for 5 months, starting at approximately 3-months of age, prior to the onset of significant Aβ pathology (Paul et al., 2018; Rosas-Hernandez et al., 2020). Additional control Tg-SwDI groups and WTs received plain drinking water only. Sample sizes were as follows: WT male n=14 (1 early death), Tg-SwDI male control n=15, telmisartan-treated male Tg-SwDI n=12 (3 early deaths), lisinopril-treated male Tg-SwDI n=11 (4 early deaths), WT female n=14 (1 early death), Tg-SwDI female control n=12, telmisartan-treated female Tg-SwDI n=13, lisinopril-treated female Tg-SwDI n=14.

#### 2.2.1. Telmisartan

Powdered drug was dissolved in 1-2 mL of 0.1 M sodium hydroxide, then mixed into reverse osmosis water, to achieve the calculated concentration of 1 mg/kg/day. pH was adjusted to 7.0-7.4 with 0.1 M hydrochloric acid. Due to light-sensitive properties of the drug (Ma et al., 2022), solutions were covered in aluminum foil at all times.

#### 2.2.2. Lisinopril

Powdered drug was dissolved in reverse osmosis water to reach the calculated concentration of 15 mg/kg/day.

### 2.3. Blood pressure (BP) measurements

The goal of treatment with RAS-inhibiting drugs was to provide doses that were neuroprotective without affecting BP. Hypertension, especially in mid-life, is a major risk factor for dementia (Lennon et al., 2024; Loera-Valencia et al., 2021; Santisteban et al., 2023). However, the Tg-SwDI strain is not hypertensive and, therefore, using drug doses that do not affect BP allowed us to avoid the confound of BP reduction and focus solely on the neuroprotective effects of the drugs. To ensure that sub-depressor doses of the drugs were being given, BP was measured 2 and 4 months after the start of the drug treatment via the tail-cuff method (Daugherty et al., 2009; Wang et al., 2017) using a BP-200 Blood Pressure Analysis System (Visitech Systems). Measurements were taken for 4 consecutive days, with day 1 being used as acclimation. Each day, 7 preliminary measurements were taken for acclimation to the procedure, followed by 10 replicate measurements that were averaged out for the daily BP of each mouse. The 4 daily averages were averaged together in the end to determine each mouse’s average BP at each time point.

### 2.4. Behavior testing

All behavior testing occurred between 10 AM and 4 PM, during the dark cycle. Before the start of each test, mice were allowed to acclimate to the room for at least 30 minutes. During testing, the test administrator stayed out of sight. Behavior testing equipment was cleaned with Peroxigard® disinfectant spray before and after each trial. A camera was suspended above the testing site to record each test/trial. Behavior videos were analyzed using either Noldus EthoVision v16 software or manual raters blinded to the mouse’s sex, genotype, and treatment condition.

#### 2.4.1. Open field (OF) test

Mice were placed into the OF chamber for 10 minutes each. Each chamber was made of white acrylic, with smooth walls and floors measuring 45 cm (length) x 45 cm (width) x 45 cm (height). The center of each chamber was defined by dividing its surface into 4 columns and 4 rows, and taking the middle 2, resulting in a center that measured 22.5 cm (length) x 22.5 cm (width) x 22.5 cm (height). Distance traveled and peripheral vs. center activity were measured.

#### 2.4.2. Novel object recognition (NOR) test

Mice were placed into the test chamber (same as OF) for a 10-minute training trial, during which they were allowed to explore two identical objects, followed by a 1-hour rest period in their home cage. After 1 hour, mice were placed back in the chambers for a 5-minute test trial, with one of the “familiar” objects being replaced by a different (“novel”) object. Time spent exploring each object was recorded for the test trials, with intact object recognition memory indicated by a preference for spending time with the novel over familiar object. Mice with total object exploration time of less than 2 seconds were excluded from the analysis (4 Tg-SwDI male controls, 1 telmisartan-treated Tg-SwDI male, 2 lisinopril-treated Tg-SwDI males, 1 WT female, 3 Tg-SwDI female controls, 1 telmisartan-treated Tg-SwDI female, and 6 lisinopril-treated Tg-SwDI females).

#### 2.4.3. Object placement test (OPT)

Mice were placed into the test chamber (same as OF and NOR) for a 10-minute training trial, during which they were allowed to explore two identical objects, side-by-side, after which they were placed back in their cages for a 24-hour retention period. After 24 hours, mice were placed back in the chambers for a 10-minute test trial, in which one of the objects was moved to a new location within the chamber (“displaced” object). Time spent exploring each object was recorded for the test trials, with intact spatial memory indicated by a preference for spending time with the displaced over familiar object. Interactions of less than 2 seconds were not considered in the analysis (6 Tg-SwDI male controls, 5 telmisartan-treated Tg-SwDI males, 3 lisinopril-treated Tg-SwDI males, 2 Tg-SwDI female controls, 3 telmisartan-treated Tg-SwDI females, and 4 lisinopril-treated Tg-SwDI females).

#### 2.4.4. Barnes maze test

The Barnes maze consisted of a raised circular platform (diameter=122 cm), which had 40 holes around its circumference, one of which (“target hole”) contained a dark “escape box” underneath. The location of the target hole and visual cues were constant for each mouse for the duration of the experiment. Due to their natural aversion to bright open spaces as prey animals, mice are motivated to find the target hole and enter the dark “escape box”. The platform was enclosed by four white curtains with visual cues on them to provide landmarks to aid in locating the target hole. Training trials were performed for 5 days, with 2 trials per day (1-hour inter-trial interval), to assess spatial learning. During each training trial, mice were allowed to explore the maze for a maximum of 3 minutes, with the goal of finding the “escape box”. Latency to find target hole (which contains the “escape box”) was recorded. For all the trials where they did not find and/or enter the box, they were guided to it by the person administering the test and allowed to stay there for approximately 1 minute. Probe trials were also performed to assess spatial memory. Approximately 24 hours after the 5 days of training trials, a probe trial was performed (1-day probe), during which mice were allowed to explore the maze for 2 minutes in the absence of the “escape box”. A second probe trial was performed in the same manner a week after the last day of training (7-day probe). During probe trials, spatial memory was measured by latency to find the target hole, first hole score (# of holes away from the target hole for the mouse’s first hole visit, with a maximum of 20), % holes visited in the target quadrant, and time spent in the target quadrant.

### 2.5. Postmortem Analyses

#### 2.5.1. Brain sectioning

Brains were cut coronally at a thickness of 20-microns using a cryostat. Sections were thaw-mounted onto microscope slides (Fisherbrand^TM^ Tissue Path Superfrost^TM^ Plus Gold Slides), starting at approximately 1.10 mm rostral to Bregma and ending at approximately −2.91 mm caudal to Bregma (Franklin & Paxinos, 1997). Sections were mounted on 3 sets of slides: one 6-slide-pre-hippocampal set, and two 8-slide-post-hippocampal sets. Sections were left to air-dry on the slides for approximately 30 minutes before being stored at −20°C until used for the immunohistochemistry protocol.

#### 2.5.2. Immunohistochemistry (IHC)

Collagen IV immunohistochemical and Thioflavin-S staining: Slides were removed from the −20 °C freezer and allowed to come to room temperature for approximately 20 minutes. The tissue sections were then fixed with 1,000 μL of paraformaldehyde (PFA;4% in PBS) for 10 minutes at room temperature followed by three 1,000 μL-washes for 3 minutes each with 1X PBS with 0.01% sodium azide. Immediately after the third wash, the tissue sections were incubated for 30 minutes at room temperature with 1,000 μL of permeabilization solution [0.3% Triton X-100 (a nonionic surfactant and emulsifier) in 1X PBS with 0.01% sodium azide]. Tissue sections were incubated for an additional 30 minutes at room temperature with 1,000 μL blocking solution (4% donkey serum in permeabilization solution) to block nonspecific antibody binding. Finally, tissue was incubated overnight at 4 °C with 325 μL of collagen IV antibody (1:200; Invitrogen, Waltham, MA, Catalog # PA1-28534, Lot # ZG4390042) for blood vessel localization. On the next day, slides were washed three times with 1,000 μL of 1X PBS with 0.01% sodium azide, for 3 minutes each, to wash off unbound primary antibody. The tissue sections were then incubated with Donkey anti-Rabbit IgG (H+L) Highly Cross-Adsorbed Secondary Antibody, Alexa Fluor^TM^ 594 (Invitrogen, Waltham, MA, Catalog # A21207, Lot # 2747441) for 1 hour at room temperature in the dark. Following the 1-hour incubation, tissue was washed 3 times with 1,000 μL of 1X PBS with 0.01% sodium azide, for 3 minutes each, at room temperature. Tissue was incubated for 15 minutes with 400 μL of 0.0125% thioflavin-S in 50% 1X PBS/50% ethanol at room temperature in the dark, to stain fibrillary Aβ. After 15 minutes, the tissue sections were washed two times with 1,200 μL of 50% 1X PBS with 0.01% sodium azide/50% ethanol, for 3 minutes each, to reduce background staining, followed by three 3-minute-washes with 1,000 μL of 1X PBS with 0.01% sodium azide. Slides were cover slipped with 100 μL of Prolong Gold mounting media, then left in the dark overnight to cure at room temperature. On the next day, the edges of cover slipped slides were sealed with clear nail polish then kept in the dark at 4 °C to be imaged in the following 1 to 3 days.

Glial fibrillary acidic protein (GFAP) immunohistochemical staining: The same protocol as described in the previous paragraph was used except for the omission of incubation with thioflavin-S solution and the subsequent wash steps with 50% 1X PBS with 0.01% sodium azide/50% ethanol. Additionally, the Prolong Gold mounting media with 4’-6-diamidino-2-phenylindole (DAPI) was used to stain nuclei. GFAP-like immunoreactivity was labeled with a Rabbit Anti-GFAP Antibody Picoband® (Boster Bio, Pleasanton, CA, Catalog # PB9082, Lot # 21BP75L15) for astrocyte localization.

Ionized calcium-binding adaptor molecule 1 (Iba1) immunohistochemical staining: Followed the same protocol as for GFAP staining using Anti-Iba1, Rabbit (for Immunocytochemistry) (FUJIFILM Wako Pure Chemical Corporation, Richmond, VA, Catalog # 019-19741, Lot # CKL4367) for microglial localization.

Imaging was done using an Olympus IX73 fluorescent microscope with cellSens® software, followed by data analysis using ImageJ software to determine % area covered by the stain or target protein/peptide immunostaining in the following regions of interest (ROIs): dentate gyrus, cornu ammonis 1 (CA1); sensorimotor cortex (SMC); ventrolateral thalamus; and dorsal subiculum. Of note, thioflavin-S staining for fibrillar amyloid was only reported in Tg-SwDI mice, as WT mice did not show measurable accumulation in the brain, as expected.

### 2.6. Statistical Analysis

Data were analyzed using 2-way ANOVAs (factors: strain/treatment and sex), followed by Tukey’s multiple comparisons post-hoc tests to determine differences between groups. One-sample t-tests were performed to compare delivered drug doses to the target dose. Outliers were identified by Grubbs’ test and removed prior to analyses. Threshold for statistical significance was set at p<0.05, while trends were described when p<0.10. Results are presented as mean±SEM. GraphPad Prism Version 10.3.1 (GraphPad Prism, San Diego, CA) was used for all data analyses and graphing.

## 3. Results

### 3.1. Neither telmisartan nor lisinopril altered systolic blood pressure (BP) of Tg-SwDI mice

Drug intake was calculated weekly for each cage in mg/kg BW/day and averaged across the study. A one-sample t-test was done to determine whether target doses were reached, comparing the average drug intake for each cage of mice and the target doses of each drug (1 mg/kg BW/day for telmisartan and 15 mg/kg BW/day for lisinopril). Statistical analysis showed that the consumed doses did not significantly differ from the target doses in either the males (p=0.333 and p=0.198 for telmisartan and lisinopril, respectively) or the females (p=0.423 and p=0.781 for telmisartan and lisinopril, respectively).

Drugs did not significantly alter systolic BP in either male or female Tg-SwDI mice compared to their untreated counterparts. Two months into the drug treatment (**Table 1)**, all Tg-SwDI groups displayed significantly lower systolic BP than WTs, when data was collapsed on sex. When separated by sex, that was still the case among the female mice. However, among the male mice, telmisartan-treated Tg-SwDI mice was the only group to display significantly lower BP than WTs. Four months into the drug treatment (**Table 1)**, lisinopril-treated Tg-SwDI mice was the only group to display significantly lower BP than WTs, when data was collapsed on sex. When separated by sex, that was only true among the female mice, while no statistically significant differences were observed among the male groups. Overall, male mice displayed higher systolic BP than females; this sex difference was maintained across all Tg-SwDI groups at 2 months, and all 4 groups at 4 months (**Table 2)**.

**Table 1.** Statistical analysis of BP 2 and 4 months after the start of drug treatment across all cohorts.

| 2-months* |  |  |  |  |  |  |
| --- | --- | --- | --- | --- | --- | --- |
|  | p-value |  |  | Significant? |  |  |
|  | Collapsed on sex | ♂ | ♀ | Collapsed on sex | ♂ | ♀ |
| WT vs. Tg-SwDI | <0.0001 | 0.4756 | <0.0001 | Yes | No | Yes |
| WT vs. Tg-SwDI TELM | <0.0001 | 0.0214 | <0.0001 | Yes | Yes | Yes |
| WT vs. Tg-SwDI LIS | <0.0001 | 0.2528 | <0.0001 | Yes | No | Yes |
| Tg-SwDI vs. Tg-SwDI TELM | 0.5577 | 0.3868 | 0.9907 | No | No | No |
| Tg-SwDI vs. Tg-SwDI LIS | 0.6031 | 0.9701 | 0.5810 | No | No | No |
| Tg-SwDI TELM vs. Tg-SwDI LIS | 0.9995 | 0.6613 | 0.7404 | No | No | No |
| 4-months** |  |  |  |  |  |  |
|  | p-value |  |  | Significant? |  |  |
|  | Collapsed on sex | ♂ | ♀ | Collapsed on sex | ♂ | ♀ |
| WT vs. Tg-SwDI | 0.4297 | 0.6684 | 0.7396 | No | No | No |
| WT vs. Tg-SwDI TELM | 0.1257 | 0.4586 | 0.3467 | No | No | No |
| WT vs. Tg-SwDI LIS | 0.0025 | 0.1145 | 0.0245 | Yes | No | Yes |
| Tg-SwDI vs. Tg-SwDI TELM | 0.8908 | 0.9775 | 0.9306 | No | No | No |
| Tg-SwDI vs. Tg-SwDI LIS | 0.1617 | 0.6242 | 0.3022 | No | No | No |
| Tg-SwDI TELM vs. Tg-SwDI LIS | 0.5327 | 0.8719 | 0.6514 | No | No | No |

Statistical significance was set at $p < 0.05$ .
\*A main effect of group was observed 2 months after the start of treatment ( $p < 0.0001$ )
\*\*A main effect of group was observed 4 months after the start treatment ( $p = 0.0049$ )

**Table 2.** Statistical analysis for sex differences in BP between each group across all cohorts.

| **2-months\*** |  |  |
| --- | --- | --- |
|  | **p-value** | **Significant?** |
| **WT ♂ vs. WT ♀** | 0.7889 | No |
| **Tg-SwDI ♂ vs. Tg-SwDI ♀** | 0.0001 | Yes |
| **Tg-SwDI TELM ♂ vs. Tg-SwDI TELM ♀** | 0.0079 | Yes |
| **Tg-SwDI LIS ♂ vs. Tg-SwDI LIS ♀** | <0.0001 | Yes |
| **4-months\*\*** |  |  |
|  | **p-value** | **Significant?** |
| **WT ♂ vs. WT ♀** | 0.0044 | Yes |
| **Tg-SwDI ♂ vs. Tg-SwDI ♀** | 0.0063 | Yes |
| **Tg-SwDI TELM ♂ vs. Tg-SwDI TELM ♀** | 0.0044 | Yes |
| **Tg-SwDI LIS ♂ vs. Tg-SwDI LIS ♀** | 0.0013 | Yes |

Statistical significance was set at $p < 0.05$ .
\*A main effect of sex was observed 2 months after the start of treatment ( $p < 0.0001$ )
\*\*A main effect of sex was observed 4 months after the start of treatment ( $p < 0.0001$ )

### 3.2. Tg-SwDI mice showed significantly less exploratory behavior than WTs, which is not improved by drug treatment

The open field test was performed to assess general locomotor activity and anxiety-like behavior. Females showed significantly higher activity levels than males, represented by distance traveled; and significantly less anxiety-like behavior, represented by increased % center time. No statistically significant sex differences were observed in rearing and grooming behavior, which are additional measurements of exploratory and anxiety-like behaviors, respectively. Similar results were observed between treated and untreated Tg-SwDI and WT mice, with Tg-SwDI being significantly less active than WTs (**Figure 2**).

**Figure 2.**
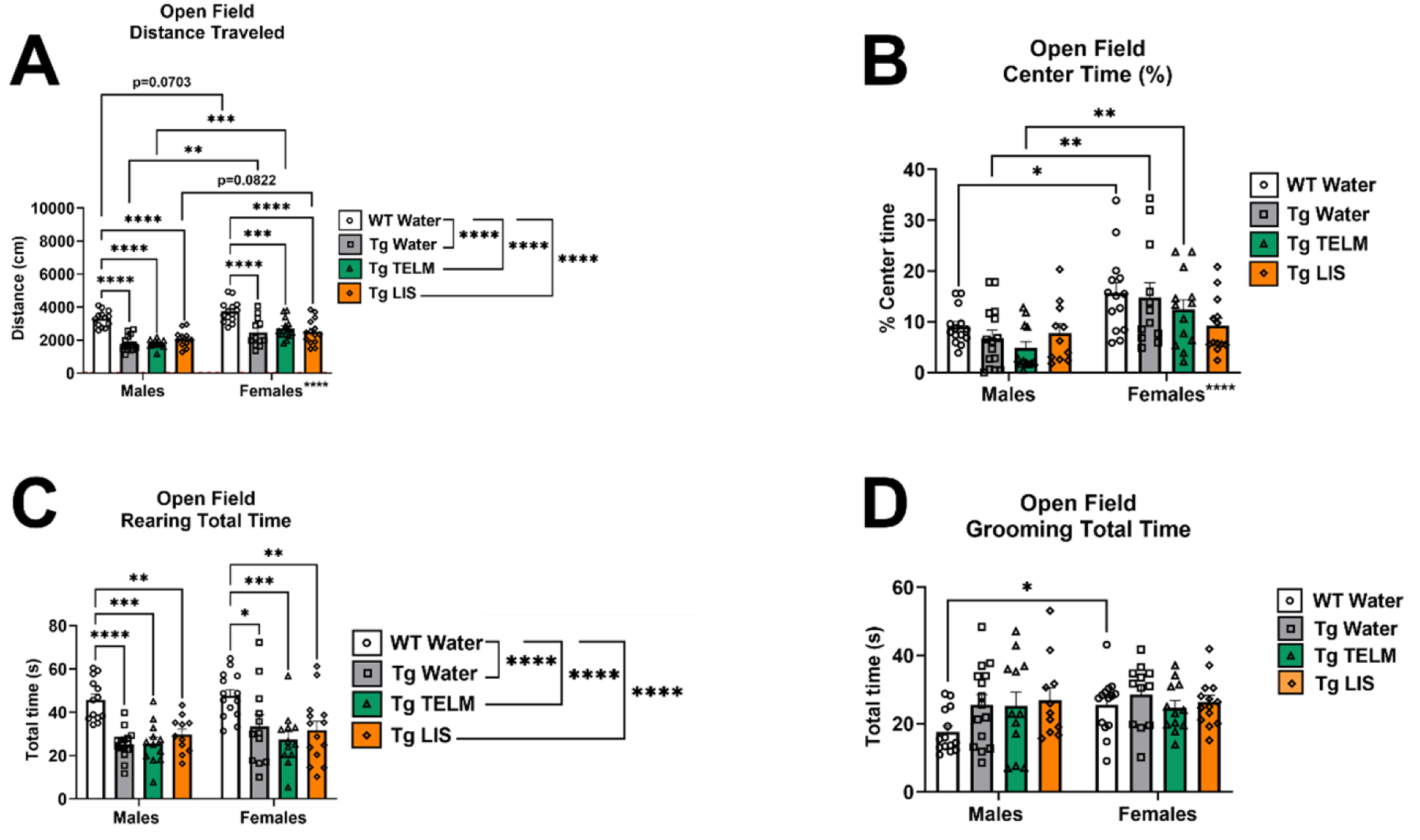
Open field data for exploratory and anxiety-like behaviors. **(A)** When collapsed on sex, data showed that Tg-SwDI mice of all groups moved significantly less than WTs (p<0.0001 for Tg-SwDI controls, p<0.0001 for telmisartan-treated Tg-SwDI mice, and p<0.0001 for lisinopril-treated Tg-SwDI mice). When separated by sex, the same was true for both sexes (p<0.0001 for all comparisons except that of WT females with telmisartan-treated females, which had p=0.0002). Females moved significantly more than males (p<0.0001), which was driven by the Tg-SwDI controls (p=0.0051) and telmisartan-treated Tg-SwDI (p=0.0002) groups. No effect of drug treatment was observed. **(B)** When collapsed on sex, data showed no significant differences between groups in time spent in the center of the arena. When separated by sex, this remained true in both males and females. No effect of drug treatment observed. Females spent significantly more time in the center of the arena compared to males (p<0.0001). However, that was only observed in WTs (p=0.0103), Tg-SwDI controls (p=0.0024), and telmisartan-treated Tg-SwDI mice (p=0.0054). **(C)** When collapsed on sex, data showed that WTs displayed significantly more rearing behavior than Tg-SwDI mice of all groups (p<0.0001 for Tg-SwDI controls, p<0.0001 for telmisartan-treated Tg-SwDI mice, and p<0.0001 for lisinopril-treated Tg-SwDI mice). The same was observed in both males and females when data was separated by sex. No effect of drug treatment or sex differences were observed. **(D)** When collapsed on sex, data showed no significant differences in grooming behavior between groups. The same was observed in both males and females when data was separated by sex. No effect of drug treatment was observed. WT females spent significantly more time grooming than WT males (p=0.0353).

### 3.3. Telmisartan rescued object recognition memory, particularly in Tg-SwDI males

The novel object recognition (NOR) test was performed to assess object recognition memory. In line with findings shown in the OF test, Tg-SwDI mice displayed less time exploring the objects than WTs, a trend that was unaffected by drug treatment, as seen with the time spent exploring both objects (**Figure 3A**). No differences were observed between treatment groups when data was separated by sex. Additionally, telmisartan rescued the impairment in object recognition memory exhibited by Tg-SwDI mice, particularly in males, while lisinopril had no effect on object recognition memory in either sex (**Figure 3B**).

**Figure 3.**
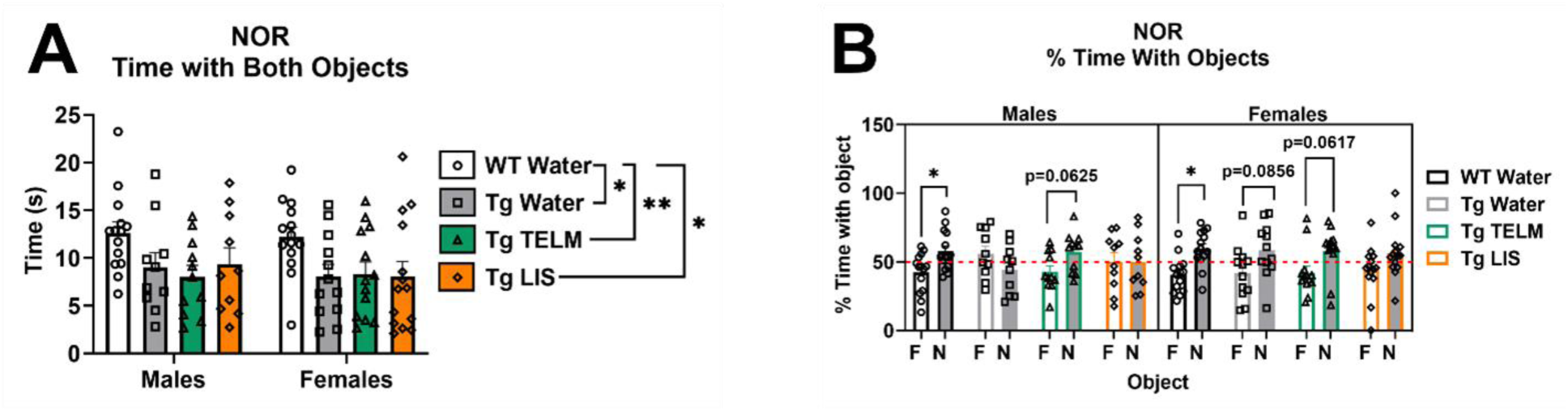
NOR data for learning and memory. **(A)** When the data was collapsed on sex, all Tg-SwDI groups spent significantly less time than WTs exploring objects (p=0.0241 for Tg-SwDI controls, p=0.0309 for telmisartan-treated Tg-SwDI mice, and p=0.0088 for lisinopril-treated Tg-SwDI mice). When separated by sex, data showed no differences between WTs and Tg-SwDI controls, or between Tg-SwDI groups in either sex. No sex differences were observed. **(B)** A paired t-test analysis was done to compare time spent with each object [novel (N) versus familiar (F)]. WT males showed a statistically significant preference for the novel object compared to the familiar (M=15.69, SD=29.09), t(13)=2.019, p=0.0323; while Tg-SwDI controls showed a non-significant preference for the familiar object (M=-11.39, SD=35.33), t(9)=1.019, p=0.1673. In the males, telmisartan showed potential in normalizing behavior, while lisinopril did not result in improvement. Similar to the males, WT females showed a statistically significant preference for the novel object compared to the familiar one (M=19.05, SD=26.96), t(13)=2.644, p=0.0101; while transgenic controls showed a non-significant preference for the novel object compared to the novel one (M=16.59, SD=39.24), t(11)=1.464, p=0.0856. Telmisartan showed potential in normalizing behavior, while lisinopril did not result in improvement.

### 3.4. No effect of drug treatment was observed on performance of male or female Tg-SwDI mice in the object placement test (OPT)

The OPT was performed to assess spatial memory. In line with results from the OF and NORT, Tg-SwDI mice showed significantly lower object exploratory behavior than WTs, as seen with the time spent exploring both objects in the OPT (p=0.0002 for Tg-SwDI controls, p<0.0001 for telmisartan-treated Tg-SwDI mice, and p=0.0010 for lisinopril-treated Tg-SwDI mice). However, when separated by sex, this was only statistically significant in males (p=0.0002 for Tg-SwDI controls, p=0.0001 for telmisartan-treated Tg-SwDI mice, and p=0.0035 for lisinopril-treated Tg-SwDI mice). No effect of drug treatment was observed in either sex, and neither were there any sex differences. Additionally, a paired t-test analysis performed to compare time spent with each object, showed no significant effect of drug treatment on performance of males or females, as no groups showed preference for the displaced object (data not shown).

### 3.5. Drug treatment showed potential benefits for preserving spatial memory in Tg-SwDI mice in the Barnes maze

The Barnes maze was performed to assess spatial learning and memory. Results from Barnes maze training trials showed that WTs were significantly faster than Tg-SwDI mice in finding the target hole, indicating an impairment of spatial learning in Tg-SwDI mice. This was observed in both sexes and was unaffected by drug treatment (**Figures 4A -B**).

**Figure 4.**
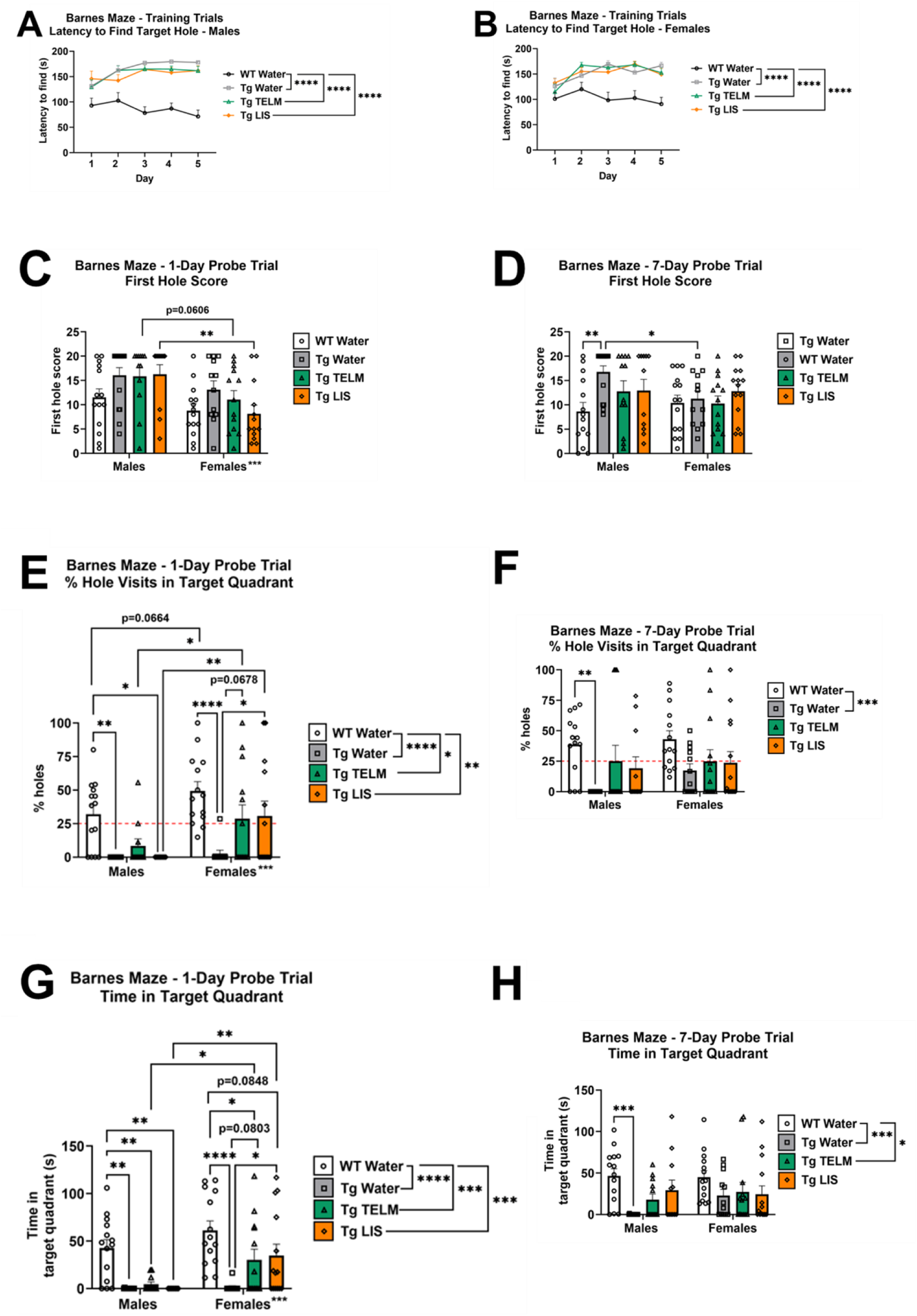
Barnes maze data for spatial learning and memory. **(A)** Latency to find the target hole was reported in the males of each group and averaged out for each day of the training trials. A significant difference was observed between WTs and all Tg-SwDI groups considering all five days of training trials (p<0.0001 for all three comparisons). WTs were the quickest and improved with time (71.5 s on day 5 versus 92.9 s on day 1), while Tg-SwDI controls were the slowest and got worse with time (178.1 s on day 5 versus 131.3 s on day 1). No effect of drug treatment was observed. **(B)** Latency to find the target hole was reported in the females of each group and averaged out for each day of the training trials. A significant difference was observed between WTs and all Tg-SwDI groups considering all five days of training trials (p<0.0001 for all three comparisons). WTs were the quickest and improved with time (90.7 s on day 5 versus 100.9 s on day 1), while Tg-SwDI controls were the slowest and got worse with time (166.4 s on day 5 versus 125.9 s on day 1). No effect of drug treatment was observed. **(C)** When collapsed on sex, data showed a significant difference between WTs and Tg-SwDI controls (p=0.0497). No effect of drug treatment was observed in the males. In the females, both drugs appeared to improve performance, but results were not significantly different from those of Tg-SwDI controls (p=0.8539 and p=0.2075 for telmisartan and lisinopril, respectively). Males of all groups performed significantly worse than the females (p=0.0003) on the 1-day probe trial; however, this was only significant between the lisinopril groups (p=0.0021). A non-parametric analysis was performed due to many mice not finding the escape hole in the time frame of the test. **(D)** When collapsed on sex, data showed a significant difference between WTs and transgenic controls (p=0.0460). In the males, Tg-SwDI controls were the only ones to perform significantly worse than the WTs (p=0.0042); while in the females, no group differences were observed. Males and females performed similarly on the 7-day probe trial, although male Tg-SwDI controls performed significantly worse than females of the same group (p=0.0262). A non-parametric analysis was performed due to many mice not finding the escape hole in the time frame of the test. **(E)** When collapsed on sex, data showed a significant difference was observed between WTs and all Tg-SwDI groups (p<0.0001 for WTs vs. Tg-SwDI controls; p=0.0097 for WTs vs. telmisartan-treated Tg-SwDI mice, and p=0.0027 for WTs vs. lisinopril-treated Tg-SwDI mice). In the males, only telmisartan showed potential in normalizing behavior (p=0.1129 compared to WTs). In the females, drug treatment with both drugs improved performance, as no significant difference from WTs was observed (p=0.1813 and p=0.2788 for telmisartan and lisinopril, respectively), and both groups performed significantly better than the transgenic controls (p=0.0678 and p=0.0400 for telmisartan and lisinopril, respectively). Females visited significantly more holes in the target quadrant than males did (p=0.0006), driven by WTs (p=0.0664) and drug-treated groups (p=0.0474 and p=0.0041 for telmisartan and lisinopril, respectively). **(F)** When collapsed on sex, data showed significant differences between WTs and Tg-SwDI controls (p=0.0005). In males, Tg-SwDI controls performed significantly worse than WTs (p=0.0031). Drug treatment improved performance and normalized behavior compared to WTs. In females, no group differences were observed. No significant differences in performance between males and females were observed on the 7-day probe. **(G)** When collapsed on sex, data showed significant differences between WTs and all Tg-SwDI groups on time spent in the target quadrant (p<0.0001 for transgenic controls, p=0.0003 for telmisartan- and lisinopril-treated Tg-SwDI mice). In the males, no effect of drug treatment was observed. In the females, lisinopril improved performance compared to the transgenic controls (p=0.0272) and normalized behavior compared to WTs (p=0.0848). While telmisartan also improved behavior, it remained significantly different from WTs (p=0.0347), but fell short of significance compared to Tg-SwDI controls (p=0.0803). Overall, females spent significantly more time in the target quadrant than males (p=0.0009), driven mainly by the treatment groups (p=0.0338 and p=0.0047 for telmisartan and lisinopril, respectively). **(H)** When collapsed on sex, data showed significant differences between WTs and Tg-SwDI controls (p=0.0006), as well as WTs and telmisartan-treated Tg-SwDI mice (p=0.0404), but not lisinopril-treated Tg-SwDI mice (p=0.1352), in the time spent in the target quadrant. In males, Tg-SwDI controls performed significantly worse than WTs (p=0.0007). Drug treatment improved performance and normalized behavior compared to WTs (p=0.0982 for telmisartan-treated Tg-SwDI mice, and p=0.5203 for lisinopril-treated Tg-SwDI mice), but fell short of significance compared to Tg-SwDI controls (p=0.4631 and p=0.0948 for telmisartan and lisinopril, respectively). In females, no group differences were observed. No significant difference in the time spent in the target quadrant on the 7-day probe trial were observed between males and females (p=0.2998).

Results from the 1-day and 7-day probe trials gave insight into shorter- and longer- term memory, respectively. Data consisted of first-hole score (how many holes from the target hole was the first hole visited) (**Figures 4C-D**), percentage of holes visited that were located in the target quadrant (**Figures 4E-F**), and time spent in the target quadrant (**Figures 4G-H**).

#### First Hole Score

In the males, no significant difference between treatment groups were observed during the 1-day probe trial. However, Tg-SwDI controls performed significantly worse than WTs during the 7-day probe, while treatment with either drug normalized the performance of the Tg-SwDI mice. In the females, no significant differences were observed between treatment groups during either probe trial. Sex differences were observed only during the 1-day probe trial, driven by the drug-treated groups, with drug-treated females performing better than drug-treated males. Overall sex differences were not observed during the 7-day probe trial; although, when separated by treatment, untreated male Tg-SwDI mice performed significantly worse than their female counterparts.

#### % Hole Visits in Target Quadrant

When collapsed on sex, Tg-SwDI mice, regardless of treatment group, performed significantly worse than WT mice during the 1-day probe trial. However, on the 7-day probe trial, treatment with either drug normalized behavior to WT levels. In the males, telmisartan partially rescued behavior (to that of WTs) on the 1-day probe trial, while both drugs achieved that on the 7-day probe trial. In the females, both drugs normalized behavior on the 1-day probe trial, while no significant differences were seen between treatment groups on the 7-day probe trial. Sex differences were observed on the 1-day probe trial (females performed better than males), though this was driven by WTs and drug-treated groups. No sex difference was observed in the 7-day probe trial.

#### Time Spent in the Target Quadrant

When collapsed on sex, Tg-SwDI mice, regardless of treatment group, performed significantly worse than WT mice during the 1-day probe trial, while lisinopril partially rescued WT behavior on the 7-day probe trial. In the males, WTs performed significantly better than all Tg-SwDI groups on the 1-day probe trial, while both drugs rescued behavior (to that of WTs) on the 7-day probe trial. In the females, both drug-treated groups performed significantly better than Tg-SwDI controls, but still significantly worse than WTs, on the 1-day probe trial. However, no significant differences were observed between treatment groups on the 7-day probe trial. Overall, results showed that treatment with both drugs had some ability to preserve cognitive function in both sexes.

### 3.6. Lisinopril partially ameliorated the deficit in vascular density in the dentate gyrus of Tg-SwDI females, while telmisartan partially ameliorated increased astrocyte density in the dentate gyrus of Tg-SwDI males

No differences in vascular density were observed in the dentate gyrus among the male mice, including drug treatment effects. However, among the females, lisinopril partially normalized the reduced vascular density of Tg-SwDI mice to that of WTs, although no significant difference was observed in comparison to untreated Tg-SwDI mice (**Figures 5A and 5C**). No drug treatment effects were observed in the Aβ pathology in either sex (**Figure 5B and 5C**).

**Figure 5.**
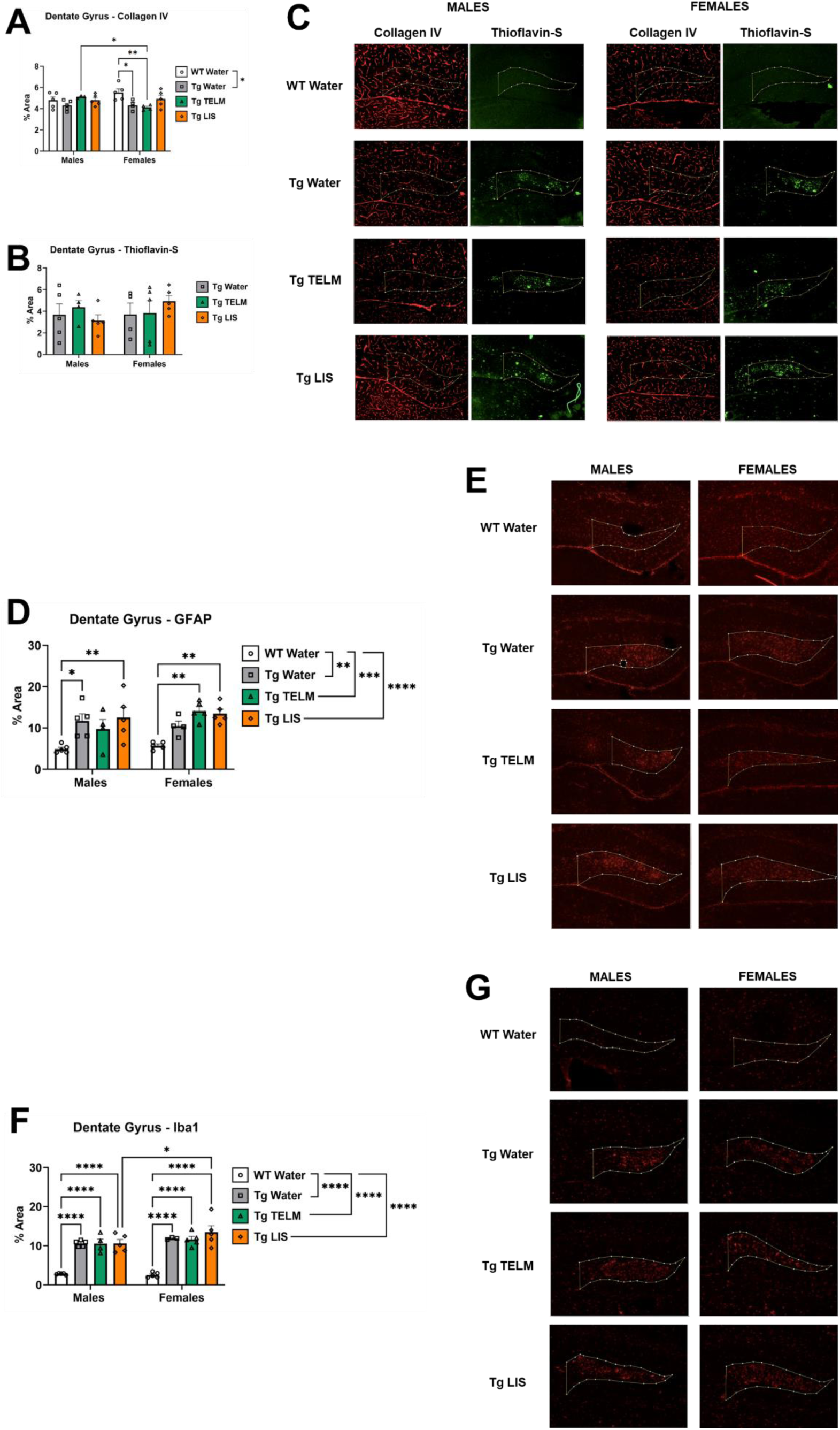
IHC – Dentate gyrus. **(A)** When collapsed on sex, Tg-SwDI controls showed significantly lower vascular density than WTs in the dentate gyrus (p=0.0203). However, when separated by sex, that was only true among the females (p=0.0242). In males, there was no effect of drug treatment on vascular density; however, in females, lisinopril partially ameliorated the deficit in vascular density in transgenic mice compared to WTs (p=0.3620), while telmisartan had no effect compared to WTs (p=0.0059). No overall sex differences were observed. However, when separated by treatment group, telmisartan-treated Tg-SwDI males presented significantly higher vascular density in the dentate gyrus than females of the same group. **(B)** When collapsed on sex, no differences on amyloid pathology were observed between treatment groups. There was no effect of drug treatment observed in the dentate gyrus in either sex. A post hoc analysis of Tg-SwDI controls versus the combined treatment groups indicated no significant differences between the groups in males or females or sex differences. Note: WTs were not included in this graph because no measurable thioflavin-S staining was observed. **(C)** Representative brain sections from 8- month-old Tg-SwDI mice stained with thioflavin-S for localization of fibrillar microvascular amyloid (green) and immunolabeled with collagen IV for identification of cerebral blood vessels (red) in the dentate gyrus. **(D)** When collapsed on sex, Tg-SwDI mice showed greater astrocyte density in the dentate gyrus than WTs (p=0.0028 for Tg-SwDI controls, p=0.0006 for telmisartan-treated Tg-SwDI mice, and p<0.0001 for lisinopril-treated Tg-SwDI mice). In the males, Tg-SwDI controls and lisinopril-treated Tg-SwDI mice were the only groups to present significantly higher astrocyte density than WTs (p=0.0116 and p=0.0040, respectively). Although telmisartan appeared to ameliorate the presence of astrocytes in the dentate gyrus of Tg-SwDI males, the decrease observed was not statistically significant, when compared to Tg-SwDI controls (p=0.8017). In the females, both drugs appeared to increase astrocyte density in the dentate gyrus, as the % area covered by astrocytes was significantly higher in both compared to WTs (p=0.0015 for telmisartan-treated Tg-SwDI females, and p=0.0035 for lisinopril-treated Tg-SwDI females). However, it was not significant when compared to the transgenic controls (p=0.3497 for telmisartan-treated Tg-SwDI females, and p=0.5176 for lisinopril-treated Tg-SwDI females). No sex differences were observed. A post hoc analysis of the transgenic controls versus the combined treatment groups indicated no significant differences between the two groups in males or females or sex differences**. (E)** Representative images for each sex and group depict the differences shown on panel D. **(F)** When collapsed on sex, Tg-SwDI mice showed significantly higher microglia density than WTs in the dentate gyrus (p<0.0001 for Tg-SwDI controls, p<0.0001 for telmisartan-treated Tg-SwDI mice, and p<0.0001 for lisinopril-treated Tg-SwDI mice). That remained true when data was separated by sex. Microglia density was increased in dentate gyrus of lisinopril-treated Tg-SwDI females compared to males (p=0.0246), although the differences compared to Tg-SwDI controls were not significant. Drug treatment did not affect microglia density in the dentate gyrus of either male or female mice. No statistically significant sex differences were observed. A post hoc analysis of the transgenic controls versus the combined treatment groups indicated no significant differences between the two groups in males or females or sex differences. **(G)** Representative images for each sex and group depict the differences shown on panel F.

In the males, telmisartan partially normalized the enhanced astrocyte density in the dentate gyrus of Tg-SwDI males, although results were not significantly different from those of Tg-SwDI controls. In the females, both Tg-SwDI drug-treated groups exhibited increased astrocyte density compared to WT controls, though the increases from untreated Tg-SwDI were not significant (**Figure 5D-E**). Additionally, Tg-SwDI mice showed significantly higher microglial density in the dentate gyrus than WTs, which was unaffected by either drug treatment in either sex (**Figures 5F-G**).

### 3.7. Lisinopril reduced microglial density in the sensorimotor cortex of Tg-SwDI females

No significant differences were observed in vascular density in the sensorimotor cortex of Tg-SwDI mice compared to WTs. Additionally, no effect of drug treatment was observed in either the vascular density, Aβ pathology, or number of Aβ plaques (**Figures 6A-D**).

**Figure 6.**
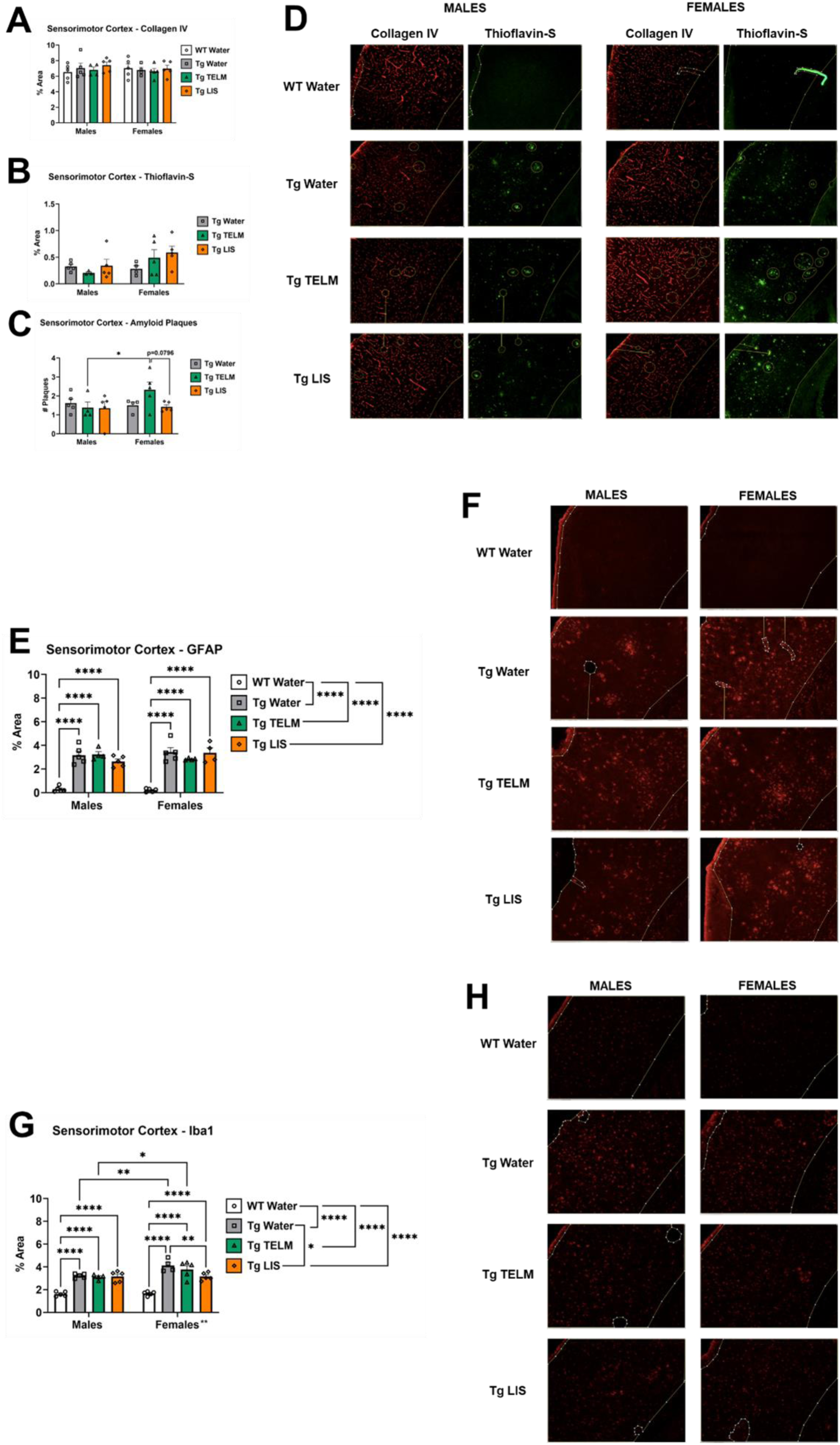
IHC – Sensorimotor cortex. **(A)** There was no statistically significant difference in vascular density between WTs and Tg-SwDI mice. Additionally, there was no effect of drug treatment in the vascular density of the sensorimotor cortex in neither males nor females. No statistically significant sex differences were observed. **(B)** There was no effect of drug treatment in Aβ pathology in the sensorimotor cortex. No statistically significant sex or treatment group differences were observed. A post hoc analysis of the Tg-SwDI controls versus the combined treatment groups indicated a trend of increased thioflavin-S with RAS-targeting drugs in the sensorimotor cortex of Tg-SwDI females (p=0.0790). Additionally, sex comparisons indicated a trend of higher levels of thioflavin-S staining in the sensorimotor cortex of drug-treated females compared to their male counterparts (p=0.0730). **(C)** No statistically significant differences in the number of Aβ plaques were observed between treatment groups. There was no effect of drug treatment in the number of Aβ plaques in the sensorimotor cortex of Tg-SwDI males. Similarly, in the females, lisinopril also had no effect. However, telmisartan-treated Tg-SwDI females presented a significantly higher number of Aβ plaques in the sensorimotor cortex, compared with males of the same group (p=0.0341). No statistically significant sex differences were observed. A post hoc analysis of the Tg-SwDI controls versus the combined treatment groups indicated no significant differences between the groups in males or females or sex differences. Note: WTs were not included in the graphs on panels B and C because no measurable thioflavin-S staining was observed. **(D)** Representative brain sections from 8-month-old Tg-SwDI mice stained with thioflavin-S for localization of fibrillar microvascular amyloid (green) and immunolabeled with collagen IV for identification of cerebral blood vessels (red) in the sensorimotor cortex. **(E)** When collapsed on sex, Tg-SwDI mice showed significantly lower astrocyte density than WTs (p<0.0001 for Tg-SwDI controls, p<0.0001 for telmisartan-treated Tg-SwDI mice, and p<0.0001 for lisinopril-treated Tg-SwDI mice). The same was observed in males and females when data was separated by sex. No effect of drug treatment or sex differences were observed. A post hoc analysis of the Tg-SwDI controls versus the combined treatment groups indicated no significant differences between the two groups in males or females or sex differences. **(F)** Representative images for each sex and group depict the differences shown on panel E. **(G)** When collapsed on sex, Tg-SwDI mice showed significantly higher microglia density than WTs (p<0.0001 for Tg-SwDI controls, p<0.0001 for telmisartan-treated Tg-SwDI mice, and p<0.0001 for lisinopril-treated Tg-SwDI mice); as well as between Tg-SwDI controls and lisinopril-treated Tg-SwDI mice (p=0.0491) Overall, Tg-SwDI females had significantly greater microglia density in the sensorimotor cortex compared to males (p=0.0050), an effect being driven mainly by the Tg-SwDI control and telmisartan-treated Tg-SwDI groups (p=0.0025 and p=0.0217, respectively). In the males, no effect of drug treatment was observed; however, in the females, lisinopril significantly reduced microglia density in the sensorimotor cortex compared to Tg-SwDI controls (p=0.0073). A post hoc analysis of Tg-SwDI controls versus the combined treatment groups indicated a significant decrease in microglial density with RAS-targeting drugs in the sensorimotor cortex of drug-treated Tg-SwDI females (p=0.0376). **(H)** Representative images for each sex and group depict the differences shown on panel G.

Tg-SwDI mice showed higher astrocyte and microglia density in the sensorimotor cortex than WTs. Drug treatment did not influence astrocyte density in either males or females (**Figures 6E-F**). While no effect of drug treatment was observed in microglia density in males, lisinopril significantly reduced it in the females, compared to Tg-SwDI controls, although it remained significantly higher than that of WTs (**Figures 6G-H**).

### 3.8. Tg-SwDI females presented greater Aβ pathology, Aβ plaque coverage, CAA, and microglia density than Tg-SwDI males in the CA1

No significant differences were observed in vascular density in the CA1 of Tg-SwDI mice compared to WTs. Additionally, no effect of drug treatment was observed in either the vascular density, Aβ pathology, % area covered by Aβ plaques, % Aβ located in blood vessels, or vessel coverage. However, sex comparisons showed that Tg-SwDI females present greater Aβ pathology, Aβ plaque coverage, and CAA than males, trends that were driven by the treatment groups across all parameters (**Figures 7A-F**).

**Figure 7.**
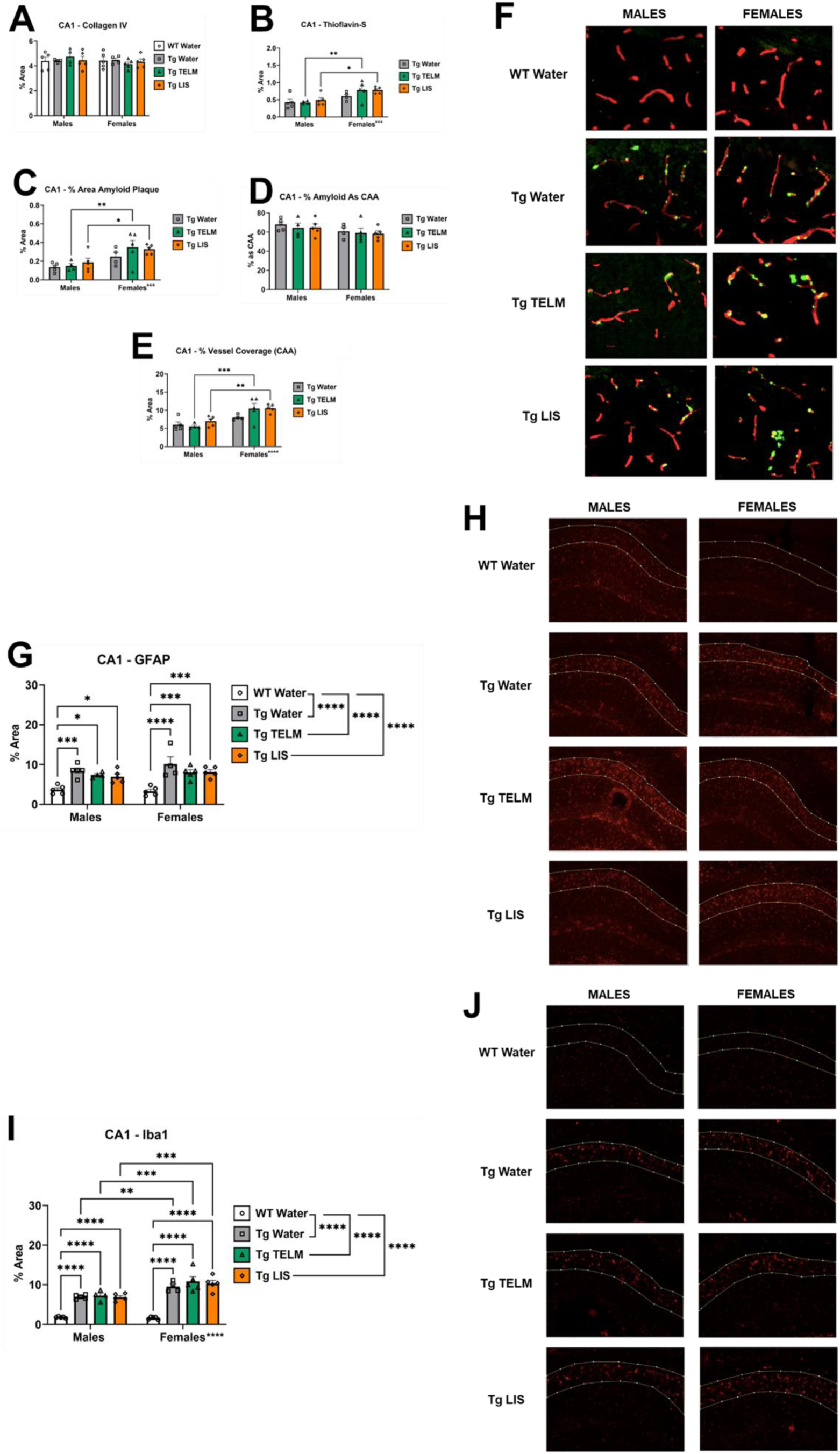
IHC – CA1. **(A)** There was no statistically significant difference in vascular density between Tg-SwDI mice and WTs. Additionally, no effect of drug treatment was observed in the vascular density of the CA1 in neither sex. No statistically significant sex differences were observed. **(B)** No statistically significant differences in Aβ distribution were observed between treatment groups. There was no effect of drug treatment in either sex. There was a statistically significant sex difference (p=0.0003) in Aβ distribution in the CA1, with Tg-SwDI females having greater Aβ distribution than males, an effect that was statistically significant between the telmisartan- and lisinopril-treated groups (p=0.0044 and p=0.0134, respectively). A post hoc analysis of Tg-SwDI controls versus the combined treatment groups indicated a trend of increased thioflavin-S with RAS-targeting drugs in the CA1 of Tg-SwDI females (p=0.0632). Additionally, sex comparisons indicated significantly higher levels of thioflavin-S staining in the CA1 of drug-treated females compared to their male counterparts (p<0.0001). **(C)** No statistically significant differences in Aβ plaque distribution were observed between treatment groups. There was no effect of drug treatment in either sex. There was a statistically significant sex difference (p=0.0005) in Aβ plaque distribution in the CA1, with Tg-SwDI females showing greater Aβ plaque coverage than males, a trend unaffected by drug treatment. However, this trend was only statistically significant between the telmisartan- and lisinopril-treated groups (p=0.0056 and p=0.0320, respectively). A post hoc analysis of the Tg-SwDI controls versus the combined treatment groups indicated no significant increase in amyloid plaque coverage in either males or females treated with RAS-targeting drugs. However, sex comparisons indicated that drug-treated females had significantly higher amyloid plaque coverage than their male counterparts (p=0.0001), while the same trend was observed between Tg-SwDI controls (p=0.0673). **(D)** There was no effect of drug treatment in the percentage of Aβ that is located on blood vessels. No statistically significant sex differences were observed. A post hoc analysis of the Tg-SwDI controls versus the combined treatment groups indicated no significant differences between the groups in males or females. **(E)** There was no statistically significant difference in vessel coverage between treatment groups. There was no effect of drug treatment in either males or females. There was a statistically significant sex difference (p<0.0001) in vessel coverage in the CA1, with Tg-SwDI females having a greater percentage of blood vessels covered by Aβ than males, a trend that was only statistically significant between the telmisartan- and lisinopril-treated groups (p=0.0004 and p=0.0038, respectively). A post hoc analysis of Tg-SwDI controls versus the combined treatment groups indicated a significant increase in vessel coverage with RAS-targeting drugs in the CA1 of Tg-SwDI females (p=0.0219). Additionally, sex comparisons indicated significantly higher levels of vessel coverage in the CA1 of drug-treated females compared to their male counterparts (p<0.0001). Note: The WT groups were not included in the graphs of panels B-E because no measurable thioflavin-S staining was observed. **(F)** Representative brain sections from 8-month-old Tg-SwDI mice stained with thioflavin-S for localization of fibrillar microvascular amyloid (green) and immunolabeled with collagen IV for identification of cerebral blood vessels (red) in the CA1. The overlap of both colors corresponds to CAA (Aβ in the blood vessel). **(G)** When collapsed on sex, Tg-SwDI mice showed significantly higher astrocyte density in the CA1 than WTs (p<0.0001 for Tg-SwDI controls, p<0.0001 for telmisartan-treated Tg-SwDI mice, and p<0.0001 for lisinopril-treated Tg-SwDI mice). Drug treatment did not affect astrocyte density in the CA1 of Tg-SwDI males or females. No sex differences were observed. A post hoc analysis of Tg-SwDI controls versus the combined treatment groups indicated no significant differences between the two groups in males or females or sex differences. However, when this data was collapsed on sex, a trend was seen between the groups, with the treatment group having lower astrocyte density than transgenic controls (p=0.0520). **(H)** Representative images for each sex and group depict the differences shown on panel G. **(I)** When collapsed on sex, Tg-SwDI mice showed significantly higher microglia density in the CA1 than WTs (p<0.0001 for Tg-SwDI controls, p<0.0001 for telmisartan-treated Tg-SwDI mice, and p<0.0001 for lisinopril-treated Tg-SwDI mice). Overall, Tg-SwDI females had significantly higher microglia density in the CA1 than Tg-SwDI males (p<0.0001), an effect that was observed in all three transgenic groups (p=0.0080 for Tg-SwDI controls, p=0.0005 for telmisartan-treated Tg-SwDI mice, and p=0.0004 for lisinopril-treated Tg-SwDI mice). No effect was drug treatment was observed in neither sex. A post hoc analysis of Tg-SwDI controls versus the combined treatment groups indicated a significant increase in microglia density with RAS-targeting drugs in the CA1 of drug-treated Tg-SwDI females compared to their male counterparts (p<0.0001). **(J)** Representative images for each sex and group depict the differences shown on panel I.

Tg-SwDI mice showed higher astrocyte and microglia density in the CA1 than WTs. Drug treatment did not influence astrocyte density in either males or females (**Figures 7G-H**). While no effect of drug treatment was observed in microglia density in males or females, Tg-SwDI females showed significantly higher density than their male counterparts (**Figures 7I-J**).

### 3.9. Lisinopril increased CAA in the ventrolateral thalamus of Tg-SwDI males

Tg-SwDI females tended to have reduced vascular density in the ventrolateral thalamus compared to WT controls; however, this was only significant in lisinopril-treated mice and approached significance in water-treated mice. No group differences in vascular density were observed in the males (**Figure 8A**). No effect of drug treatment was observed in Aβ pathology, % area covered by Aβ plaques, or in % Aβ located in blood vessels, in either sex (**Figures 8B-D**). While drug treatment had no effect on CAA in females, lisinopril significantly increased it in males (**Figure 8E**).

**Figure 8.**
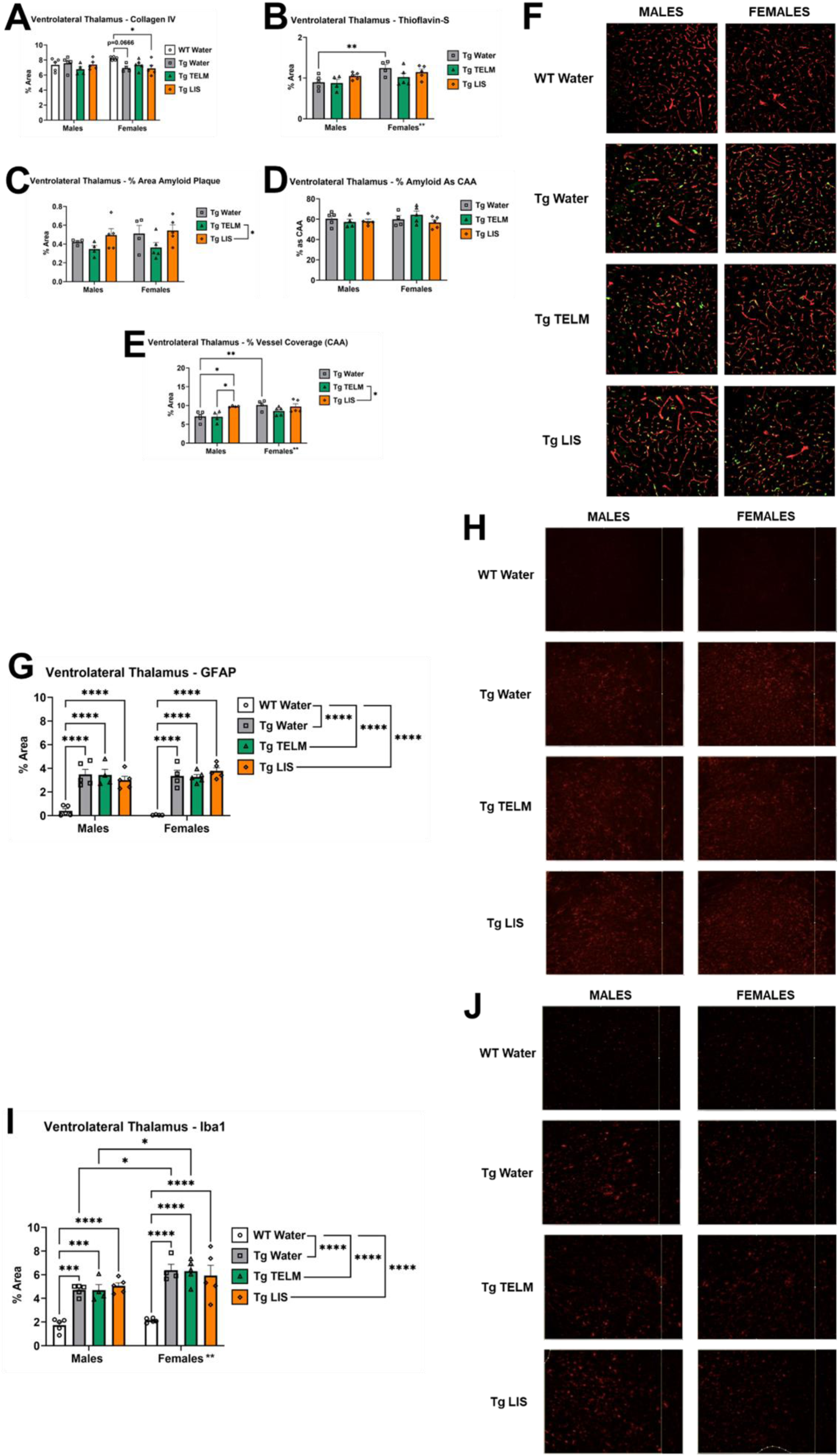
IHC – Ventrolateral thalamus. **(A)** There was no statistically significant difference in vascular density between treatment groups. In males, no effect of drug treatment was observed. In females, Tg-SwDI controls showed a slight deficit in vascular density in the ventrolateral thalamus compared to the WTs, which was further aggravated by the lisinopril treatment (p=0.0379). Additionally, no sex differences were observed. **(B)** When collapsed on sex, no statistically significant difference in Aβ pathology was observed between treatment groups. No effect of drug treatment was observed in either sex. Overall, females showed greater amyloid pathology in the ventrolateral thalamus than the males (p=0.0062). However, multiple comparison tests indicated that this was only statistically significant between the transgenic controls (p=0.0061). A post hoc analysis of T-SwDI controls versus the combined treatment groups indicated no significant differences between the groups in males or females. **(C)** Compared to telmisartan-treated mice, lisinopril-treated mice had greater Aβ plaque coverage (p=0.0309), although neither were significantly different from Tg-SwDI controls. When data was separated by sex, there was no effect of drug treatment. No statistically significant sex difference was observed. A post hoc analysis of Tg-SwDI controls versus the combined treatment groups indicated no significant differences between the groups in males or females. **(D)** There was no effect of drug treatment on the percentage of Aβ that is located on blood vessels. No statistically significant sex differences were observed. A post hoc analysis of Tg-SwDI controls versus the combined treatment groups indicated no significant differences between the groups in males or females. **(E)** Compared to telmisartan-treated mice, lisinopril-treated mice had greater vessel coverage (p=0.0126), although neither were significantly different from Tg-SwDI controls. While drug treatment did not significantly alter vessel coverage in the ventrolateral thalamus in females compared to Tg-SwDI controls, lisinopril increased % vessel coverage (CAA) in males (p=0.0192). Overall, female Tg-SwDI controls had a greater percentage of blood vessels covered by Aβ in the ventrolateral thalamus than males of the same group (p=0.0065). A post hoc analysis of Tg-SwDI controls versus the combined treatment groups indicated no significant differences between the groups in males or females. Note: WTs were not included in the graphs of panels B-E because no measurable thioflavin-S staining was observed. **(F)** Representative brain sections from 8-month-old Tg-SwDI mice stained with thioflavin-S for localization of fibrillar microvascular amyloid (green) and immunolabeled with collagen IV for identification of cerebral blood vessels (red) in the ventrolateral thalamus. The overlap of both colors corresponds to CAA (Aβ in the blood vessel). **(G)** When collapsed on sex, Tg-SwDI mice showed higher astrocyte density in the ventrolateral thalamus than WTs (p<0.0001 for Tg-SwDI controls, p<0.0001 for telmisartan-treated Tg-SwDI mice, and p<0.0001 for lisinopril-treated Tg-SwDI mice). Drug treatment did not affect astrocyte density in the ventrolateral thalamus of Tg-SwDI males or females. No sex differences were observed. A post hoc analysis of Tg-SwDI controls versus the combined treatment groups indicated no significant differences between the two groups in males or females or sex differences. **(H)** Representative images for each sex and group depict the differences shown on panel G. **(I)** When collapsed on sex, Tg-SwDI showed higher microglia density in the ventrolateral thalamus than WTs (p<0.0001 for transgenic controls, p<0.0001 for telmisartan-treated mice, and p<0.0001 for lisinopril-treated mice) (*N*=38; no outliers). Overall, Tg-SwDI females had significantly greater microglia density in the ventrolateral thalamus compared to males (p=0.0012), an effect driven mainly by the Tg-SwDI control and telmisartan-treated groups (p=0.0154 and p=0.0196, respectively). No effect of drug treatment was observed in males or females. A post hoc analysis of Tg-SwDI controls versus the combined treatment groups indicated a significant increase in microglial density with RAS-targeting drugs in the ventrolateral thalamus of drug-treated Tg-SwDI females compared to their male counterparts (p=0.0090). **(J)** Representative images for each sex and group depict the differences shown on panel I.

Tg-SwDI mice showed higher astrocyte and microglia density in the ventrolateral thalamus than WTs. Drug treatment did not reduce astrocyte density in either males or females (**Figures 8G-H**). While no effect of drug treatment was observed in microglia density in males or females, Tg-SwDI control females and telmisartan-treated Tg-SwDI females showed significantly higher density than their male counterparts (**Figures 8I-J**).

### 3.10. Lisinopril increased Aβ pathology, plaque coverage, and CAA in the dorsal subiculum of Tg-SwDI females, while telmisartan reduced microglia density in the dorsal subiculum of Tg-SwDI females

Overall, Tg-SwDI mice, regardless of drug treatment, showed increased vascular density in the dorsal subiculum compared to WT controls (**Figure 9A**); however, this was only significant in lisinopril-treated males and approached significance in telmisartan-treated males. While there was no significant effect of drug treatment within either sex for Aβ pathology, plaque coverage, % Aβ located in blood vessels, and CAA, lisinopril-treated females displayed increased Aβ pathology, plaque coverage, and CAA in the dorsal subiculum compared to their male counterparts, in addition to a reduction in % amyloid as CAA (**Figures 9B-E**).

**Figure 9.**
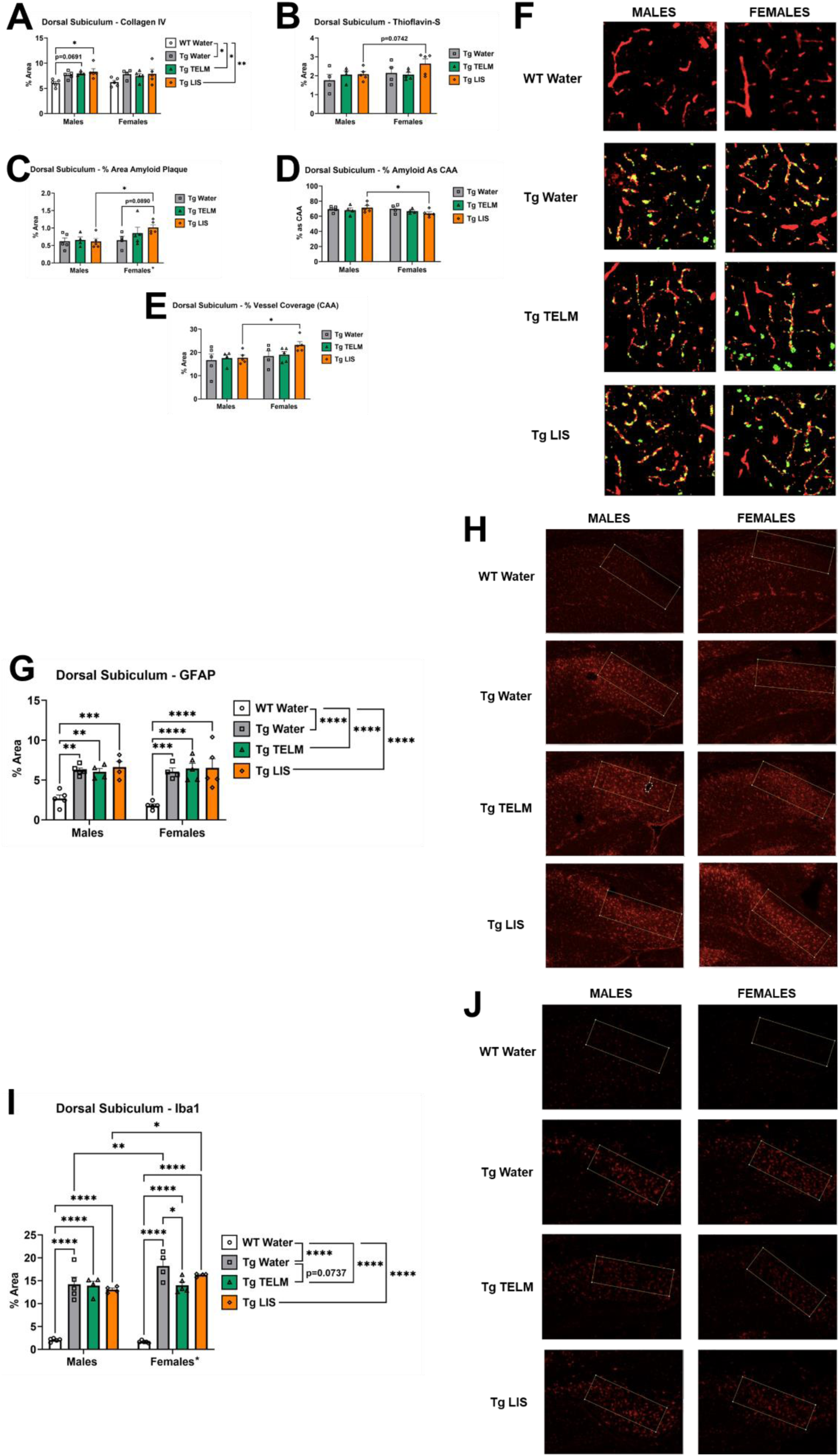
IHC – Dorsal subiculum. **(A)** When collapsed on sex, Tg-SwDI mice showed significantly higher vascular density in the dorsal subiculum than WTs (p=0.0191 for Tg-SwDI controls, p=0.0218 for telmisartan-treated Tg-SwDI mice, and p=0.0025 for lisinopril-treated Tg-SwDI mice). When separated by sex, that was only statistically significant in lisinopril-treated males (p=0.0155). No effect of drug treatment was observed in Tg-SwDI females. No sex differences were observed. **(B)** When collapsed on sex, no drug treatment differences in amyloid pathology were observed. Amyloid pathology tended to be increased in lisinopril-treated females compared to males (p=0.0742). No effect of drug treatment was observed in males or females. No overall sex differences were observed. A post hoc analysis of Tg-SwDI controls versus the combined treatment groups indicated no significant differences between the groups in males or females, or sex differences. **(C)** When collapsed on sex, no drug treatment differences in plaque coverage were observed. While no effect of drug treatment was observed in the males, a trend was observed in the females between Tg-SwDI controls and lisinopril-treated Tg-SwDI mice, with the latter showing increased plaque coverage. Females had significantly higher plaque coverage than males (p=0.0325); however, this was mainly driven by the lisinopril-treated groups. A post hoc analysis of Tg-SwDI controls versus the combined treatment groups indicated a trend of increased amyloid plaque coverage with RAS-targeting drugs in the dorsal subiculum of Tg-SwDI females (p=0.0562). Additionally, sex comparisons indicated significantly higher levels of amyloid plaque coverage in the dorsal subiculum of drug-treated females compared to their male counterparts (p=0.0108). **(D)** When collapsed on sex, no drug treatment differences were observed. The percentage of Aβ that is located on blood vessels in the dorsal subiculum was decreased in lisinopril-treated Tg-SwDI females compared to males of the same group (p=0.0260). A post hoc analysis of Tg-SwDI controls versus the combined treatment groups indicated a trend of increased Aβ levels in blood vessels with RAS-targeting drugs in the dorsal subiculum of drug-treated Tg-SwDI males compared to their female counterparts (p=0.0516). **(E)** When collapsed on sex, no drug treatment differences in vessel coverage were observed Despite lisinopril-treated Tg-SwDI females showing a significant decrease in the percentage of Aβ that is located on blood vessels in the dorsal subiculum, CAA (percentage of blood vessels covered by Aβ) was significantly increased in this group, compared to males (p=0.0369). However, this difference was not significant when compared to Tg-SwDI controls. A post hoc analysis of Tg-SwDI controls versus the combined treatment groups indicated a trend of increased vessel coverage with RAS-targeting drugs in the dorsal subiculum of drug-treated Tg-SwDI females compared to their male counterparts (p=0.0715). Note: WTs were not included in the graphs of panels B-E because no measurable thioflavin-S staining was observed. **(F)** Representative brain sections from 8-month-old Tg-SwDI mice stained with thioflavin-S for localization of fibrillar microvascular amyloid (green) and immunolabeled with collagen IV for identification of cerebral blood vessels (red) in the dorsal subiculum. The overlap of both colors corresponds to CAA (Aβ in the blood vessel). **(G)** When collapsed on sex, Tg-SwDI mice showed significantly higher astrocyte density in the dorsal subiculum than WTs (p<0.0001 for Tg-SwDI controls, p<0.0001 for telmisartan-treated Tg-SwDI mice, and p<0.0001 for lisinopril-treated Tg-SwDI mice). Drug treatment did not affect astrocyte density in the dorsal subiculum of Tg-SwDI males or females. No sex differences were observed. A post hoc analysis of Tg-SwDI controls versus the combined treatment groups indicated no significant differences between the two groups in males or females or sex differences. **(H)** Representative images for each sex and group depict the differences shown on panel G. **(I)** When collapsed on sex, Tg-SwDI mice showed significantly higher microglia density in the dorsal subiculum than WTs (p<0.0001 for Tg-SwDI controls, p<0.0001 for telmisartan-treated Tg-SwDI mice, and p<0.0001 for lisinopril-treated Tg-SwDI mice). Overall, transgenic females had significantly greater microglia density in the dorsal subiculum compared to males (p=0.0109), a trend driven mainly by the Tg-SwDI control and lisinopril-treated Tg-SwDI groups (p=0.0037 and p=0.0234, respectively). In the males, no effect of drug treatment was observed; however, in the females, telmisartan significantly reduced microglia density in the dorsal subiculum compared to Tg-SwDI controls (p=0.0118). A post hoc analysis of Tg-SwDI controls versus the combined treatment groups indicated a significant decrease in microglial density with RAS-targeting drugs in the dorsal subiculum of drug-treated Tg-SwDI females (p=0.0257). **(J)** Representative images for each sex and group depict the differences shown on panel I.

Tg-SwDI mice showed higher astrocyte and microglia density in the dorsal subiculum than WTs. Drug treatment did not affect astrocyte density in either males or females (**Figures 9G-H**). While no effect of drug treatment was observed in microglia density in the dorsal subiculum of Tg-SwDI males, telmisartan significantly reduced microglia density in the dorsal subiculum of Tg-SwDI females compared to untreated Tg-SwDI mice (**Figures 9I-J**).

## 4. Discussion

It has been established that the brain RAS is dysregulated in neurodegenerative diseases (Arregui et al., 1982; Chen et al., 2023; Cosarderelioglu et al., 2022; Miners et al., 2008; Savaskan et al., 2001), and that RAS-targeting drugs, particularly those capable of crossing the BBB, reduce dementia risk, independent of their anti-hypertensive properties (Cosarderelioglu et al., 2020; Dahlof et al., 2002; Rea et al., 2024; Ribeiro et al., 2020; Tsukuda et al., 2009; Yasar et al., 2013). This was the first study to investigate whether these drugs can prevent CAA-associated cognitive decline and neuropathology in CAA-prone, Tg-SwDI mice. Mice were treated for 5 months with sub-depressor doses of either telmisartan or lisinopril, starting at 3 months of age, followed by behavior testing at 7 months of age. Postmortem, histochemical analysis assessed the neuropathological differences between treated and untreated Tg-SwDI mice, as well as WT mice. RAS inhibitors improved some cognitive functions in Tg-SwDI mice despite no changes in Aβ burden and limited changes to vascular density and gliosis, with effects varying by sex, brain region, and behavioral task.

### 4.1. Telmisartan and lisinopril treatment during early-stage-disease showed moderate benefits for preserving some cognitive functions in Tg-SwDI mice

Tg-SwDI mice displayed significantly less exploratory behavior than WTs, as evidenced by reduced horizontal and vertical locomotor activity in the open field test, as well as diminished object exploration in the novel object recognition and object placement tasks. While reduced exploration is often interpreted as a potential marker of anxiety, this explanation is not strongly supported here, given the lack of differences in center time and grooming behavior in the open field. Alternatively, this observed behavior pattern could reflect underlying cognitive impairment, specifically deficits in perceptual speed. Perceptual speed refers to the ability to quickly and accurately detect, process, and respond to visual or other sensory information, and is considered a core cognitive domain affected in adults with CAA (Arvanitakis et al., 2011; Boyle et al., 2015). In this context, the Tg-SwDI mice’s blunted exploratory behavior suggests a diminished capacity to perceive and engage with novel environmental cues, consistent with slowed perceptual processing. Notably, this pattern of behavior was evident across multiple behavioral tasks and was not significantly altered by either telmisartan or lisinopril treatment, indicating that the drugs did not rescue impairments in sensory-cognitive integration and/or exploratory drive.

In the NOR test, mice typically show preference for the novel object (Lueptow, 2017). This behavior was observed in both sexes of WTs, while in untreated Tg-SwDI controls, a trend towards a preference for the novel object was observed in females only. Telmisartan partially restored object recognition memory in both sexes, which is in accordance with previous findings in spontaneously hypertensive rats (SHR), commonly used as a rodent model of vascular cognitive impairment and dementia. The ARB losartan rescued short-term recognition memory in the NOR test in middle-aged SHR (Tchekalarova et al., 2024).

In the Barnes maze training trials, Tg-SwDI mice were significantly slower at finding the “escape box” than WTs. Moreover, on successive training trials, the Tg-SwDI mice often failed to seek out the “escape box”, something that was not improved by drug treatment. The progressively worse performance of the Tg-SwDI mice may reflect exacerbated anxiety related to the task (which could manifest as freezing behavior) or CAA-related cognitive impairment (perceptual slowing). This test did not measure anxiety-like behavior; however, the lack of anxiety-like behavior displayed in the open field leads us to conclude that cognitive impairment played a bigger role than anxiety in the poor performance of Tg-SwDI mice in the Barnes maze training trials. Notably, when spatial memory was assessed during Barnes maze probe trials, benefits of RAS-targeting drug treatments were apparent. Treatment with either telmisartan or lisinopril improved shorter-term spatial memory (1-day probe trial) in Tg-SwDI females; however, cognitive benefits of these drugs were less obvious on the 7-day probe trial assessing longer-term spatial memory. Conversely, treatment with either telmisartan or lisinopril showed great potential for preserving cognitive function in Tg-SwDI males, significantly improving performance in both the short-term and long-term spatial memory tasks. This is in agreement with previous studies demonstrating that RAS-targeting drugs preserve spatial memory in rodent models of AD and cerebrovascular disease (Gao et al., 2018; Haraguchi et al., 2010; Kuber et al., 2023; Shindo et al., 2012).

### 4.2. Telmisartan and lisinopril had mixed effects in the vasculopathy and neuropathology of Tg-SwDI mice

Histochemistry was performed to investigate potential mechanisms of neuroprotection conferred by RAS-targeting drugs, including vascular density, Aβ distribution, and astrocyte and microglial density, in 5 brain regions: dentate gyrus [involved in learning, memory, and mood regulation (Borzello et al., 2023; Sun et al., 2023)], CA1 of the hippocampus [involved in memory consolidation (Bartsch et al., 2011)], somatosensory cortex [involved in integrating sensory information with motor function (Sessle, 2016)], ventrolateral thalamus [relay station for sensory and motor signals between the brain and the rest of the body (Torrico & Munakomi, 2025)], and dorsal subiculum [involved in memory and spatial navigation (Kitanishi et al., 2021; O’Mara, 2005)].

In Tg-SwDI mice, a significant reduction in vascular density was observed in the dentate gyrus, particularly among females; this deficit was partially ameliorated by treatment with lisinopril. Additionally, female Tg-SwDI mice exhibited reduced vascular density in the ventrolateral thalamus, which was partially rescued following treatment with telmisartan. In contrast, both male and female Tg-SwDI mice tended to show increased vascular density in the dorsal subiculum, which was not mitigated by either drug treatment. Previously, Tg-SwDI mice of unspecified sex presented decreased vascular density in the cortex, hippocampus, and thalamus (Miao et al., 2005). Other studies have looked at vascular density in animal models with parenchymal Aβ accumulation, reporting mixed results, ranging from nonsignificant changes (Delafontaine-Martel et al., 2018; Nikolajsen et al., 2015) to decreased vascular density (Fisher et al., 2022) to transient increases followed by decreases as disease progresses (Giuliani et al., 2019; Xu et al., 2020) to increased vascular density (Bennett et al., 2018). Regions reported to be most affected by abnormal vascular density in these animal models are the cortex and the hippocampus (Fisher et al., 2022; Xu et al., 2020). However, the absence of subregional analysis within the hippocampus and thalamus in the referenced studies may contribute to the discrepancies observed between their findings and ours.

Vascular density changes are typically region- and disease stage-specific (Fisher et al., 2022). Transgenic murine models with Aβ pathology have shown that vascular density is typically reduced compared to WTs, although no significant changes have also been observed, as has been summarized by Fisher et al. (Fisher et al., 2022). However, studies have documented an early, transient increase in vascular density that diminishes with disease progression, indicative of a brief angiogenic response during the initial stages (Giuliani et al., 2019; Xu et al., 2020). Chronic inflammation is often associated with angiogenesis (Whiteford et al., 2016); additionally, the reduced cerebral blood flow that results from Aβ accumulation and subsequent vessel narrowing (Thal et al., 2009) may trigger a compensatory angiogenic response (Kara et al., 2012; Zlokovic, 2011). These mechanisms could contribute to the increased vascular density observed in the dorsal subiculum of Tg-SwDI mice compared to WTs. Additionally, it has been reported that RAS-targeting drugs can normalize cerebrovascular structure in spontaneously hypertensive rats (Naessens et al., 2023), and both ACE inhibitors and ARBs are capable of promoting angiogenesis (Battegay et al., 2007). Some of our data aligns with these findings. For example, in Tg-SwDI females, telmisartan normalized the reduced vascular density in the ventrolateral thalamus.

Sex differences in Aβ pathology were apparent (females > males). Female Tg-SwDI mice had higher levels of fibrillary Aβ than males in the CA1 and ventrolateral thalamus, with smaller increases in the subiculum, while no significant sex differences were observed in the dentate gyrus or somatosensory cortex. Additionally, CAA was higher in females than males, but only the ventrolateral thalamus showed a statistically significant increase. This is in agreement with previous studies in Tg-SwDI mice and other AD mouse models (Hu et al., 2023; Maniskas et al., 2021), as well as reports that women have more severe pathology and are at greater risk for AD than men (Barnes et al., 2005; Filon et al., 2016; Koran et al., 2017; Nebel et al., 2018).

In contrast to previous studies reporting that RAS-targeting drugs attenuate Aβ (Ababei et al., 2023; Gebre et al., 2018; Li et al., 2012), neither telmisartan nor lisinopril reduced fibrillar amyloid accumulation in vascular (as CAA) or parenchymal (as plaques) compartments. In fact, Aβ pathology was modestly increased by lisinopril in the thalamus of males (CAA) and a trend in dorsal subiculum in females (plaque). We have previously shown that other interventions, including aerobic exercise (Robison et al., 2019) and environmental enrichment (Robison et al., 2020), exert cognitive-behavioral benefits despite an increase in insoluble Aβ in Tg-SwDI mice. Similar observations were made in AD mouse models (Jankowsky et al., 2005). These observations suggest that interventions, including RAS-targeting drugs, may exert neuroprotective effects through mechanisms independent of amyloid clearance, possibly through modulating neurovascular health, neuroinflammation, oxidative stress, or synaptic plasticity.

Microglia and astrocytes play critical roles in the pathogenesis of Alzheimer’s disease (AD), particularly in response to Aβ accumulation. Microglia are key immune cells in the brain that initially clear Aβ and trigger both inflammatory and regenerative responses. However, as disease progresses, they shift to a pro-inflammatory state, reducing their ability to clear Aβ and exacerbating neuronal damage (Merlo et al., 2020; Xu et al., 2023). Similarly, astrocytes, which support neuronal function, undergo reactive gliosis in response to Aβ, initially protecting neurons but ultimately contributing to neuroinflammation, oxidative stress, and impaired Aβ clearance, accelerating disease progression (Siracusa et al., 2019; Taylor et al., 2020).

Given the demonstrated anti-inflammatory effects of RAS-targeting drugs (Cantero-Navarro et al., 2021; Saavedra, 2021; Villapol et al., 2023), microglia (Iba1) and astrocyte (GFAP) coverage were assessed as measures of neuroinflammation. In line with previous findings (Miao et al., 2005; Rodriguez-Lopez et al., 2023; Xu et al., 2007), Tg-SwDI mice exhibited an increase in both microgliosis and astrogliosis across all brain regions measured (cortex, ventrolateral thalamus, dentate gyrus, dorsal subiculum, CA1), with increased severity of microgliosis in females compared to males in all regions except the dentate gyrus. Drug treatment had modest benefits for attenuating neuroinflammation, exemplified by partial rescues of astrogliosis in the dentate by telmisartan in males, microgliosis by lisinopril in the sensorimotor cortex of females, and microgliosis by telmisartan in the dorsal subiculum of females.

In our study, microglia levels were significantly elevated in females compared to males, possibly in response to the higher levels of Aβ present in the CA1 and ventrolateral thalamus of female Tg-SwDI mice. Additionally, studies have suggested that sex differences observed in microglial cells throughout life may play a causal role in the incidence and pathology of neurodegenerative diseases (Delage et al., 2021). During adulthood, females have more active microglia than males, increasing susceptibility to inflammatory conditions in the brain. Genetic factors also play a role, with the X chromosome containing more immune-related genes and epigenetic modifiers (Bobotis et al., 2023; Kodama & Gan, 2019). Finally, sex differences can also be mediated by hormones and environmental influences (Han et al., 2021). Overall, our findings are consistent with human data, with females showing greater Aβ pathology and microglial coverage than males (Casaletto et al., 2022; Oveisgharan et al., 2018; Vila-Castelar et al., 2024). Additionally, we were able to recapitulate the universal increase in astrocytes and microglia observed in the Tg-SwDI mice.

In summary, early-stage-disease interventions with RAS-targeting drugs had modest effects in preserving some cognitive functions and attenuating neuroinflammation in Tg-SwDI mice. Telmisartan partially restored object recognition memory in both sexes in the NOR test, improved short-term memory in both sexes and long-term memory in males in the Barnes maze, and partially rescued astrogliosis in the dentate gyrus of males. Lisinopril improved short-term memory in both sexes and long-term memory in males in the Barnes maze, and partially rescued microgliosis in the sensorimotor cortex and dorsal subiculum of females. It is important to note that telmisartan exerts neuroprotective effects not only through AT_1_R blockade, but also due to its partial agonist activity at peroxisome proliferator-activated receptor gamma (PPARγ) (Benson et al., 2004; Mogi et al., 2008; Pang et al., 2014). Whether the benefits observed with telmisartan in this study were enhanced by its PPARγ agonist activity remains unclear and would require future investigations involving a combination of telmisartan and a PPARγ antagonist. Additionally, the acute effects of telmisartan and lisinopril treatments during behavior testing were not evaluated in this study. Furthermore, it would be valuable to investigate whether these RAS-targeting drugs could provide protective effects in later stages of CAA.

## Supporting information

Graphical Abstract

Graphical Abstract - Publication License

Figure 1 - Publication License

Glossary

## Acknowledgments

The authors would like to thank Iliana Uribe, Kaitlin Martin, Eleanor Wind, and Valeria Cohen for their assistance with mouse care and data analysis.

## Data Statement

The data that supports the findings of this study have been deposited in Harvard’s Dataverse repository and can be found at https://dataverse.harvard.edu/dataverse/LisaSRobison-RASinCAA.

## Author Contributions

**Natalia Motzko Noto:** Data curation, Formal analysis, Investigation, Visualization, Writing – Original draft preparation, Writing – Reviewing and Editing; **Yazmin M. Restrepo:** Investigation, Writing – Original draft preparation, Writing – Reviewing and Editing; **Ariana Hernandez:** Investigation, Writing – Original draft preparation, Writing – Reviewing and Editing; **Chana Vogel:** Investigation, Writing – Original draft preparation, Writing – Reviewing and Editing; **Victoria Pulido-Correa:** Investigation, Writing – Original draft preparation, Writing – Reviewing and Editing; **Shay Moen:** Investigation, Writing – Original draft preparation, Writing – Reviewing and Editing; **Julianna Bonetti:** Investigation, Writing – Original draft preparation, Writing – Reviewing and Editing; **Vedika Chiduruppa:** Investigation, Writing – Original draft preparation, Writing – Reviewing and Editing; **Robert C. Speth:** Conceptualization, Methodology, Resources, Supervision, Validation, Visualization, Writing – Original draft preparation, Writing – Reviewing and Editing; **Lisa S. Robison:** Conceptualization, Data curation, Formal analysis, Funding acquisition, Investigation, Methodology, Project administration, Resources, Software, Supervision, Validation, Visualization, Writing – Original draft preparation, Writing – Reviewing and Editing

## References

Ababei, D. C., Bild, V., Macadan, I., Vasincu, A., Rusu, R. N., Blaj, M., Stanciu, G. D., Lefter, R. M., & Bild, W. (2023). Therapeutic Implications of Renin-Angiotensin System Modulators in Alzheimer’s Dementia. Pharmaceutics, 15(9). 10.3390/pharmaceutics15092290

Agarwal, A., Gupta, V., Brahmbhatt, P., Desai, A., Vibhute, P., Joseph-Mathurin, N., & Bathla, G. (2023). Amyloid-related Imaging Abnormalities in Alzheimer Disease Treated with Anti-Amyloid-β Therapy. Radiographics, 43(9), e230009. 10.1148/rg.230009

Arregui, A., Perry, E. K., Rossor, M., & Tomlinson, B. E. (1982). Angiotensin converting enzyme in Alzheimer’s disease: increased activity in caudate nucleus and cortical areas. J.Neurochem., 38, 1490–1492. (IN FILE)

Arvanitakis, Z., Leurgans, S. E., Wang, Z., Wilson, R. S., Bennett, D. A., & Schneider, J. A. (2011). Cerebral amyloid angiopathy pathology and cognitive domains in older persons. Ann Neurol, 69(2), 320–327. 10.1002/ana.22112

Barnes, L. L., Wilson, R. S., Bienias, J. L., Schneider, J. A., Evans, D. A., & Bennett, D. A. (2005). Sex Differences in the Clinical Manifestations of Alzheimer Disease Pathology. Arch Gen Psychiatry, 62(6), 685–691. 10.1001/archpsyc.62.6.685

Bartsch, T., Döhring, J., Rohr, A., Jansen, O., & Deuschl, G. (2011). CA1 neurons in the human hippocampus are critical for autobiographical memory, mental time travel, and autonoetic consciousness. Proceedings of the National Academy of Sciences, 108(42), 17562–17567. doi:10.1073/pnas.1110266108

Battegay, E. J., de Miguel, L. S., Petrimpol, M., & Humar, R. (2007). Effects of anti-hypertensive drugs on vessel rarefaction. Curr Opin Pharmacol, 7(2), 151–157. 10.1016/j.coph.2006.09.007

Bennett, R. E., Robbins, A. B., Hu, M., Cao, X., Betensky, R. A., Clark, T., Das, S., & Hyman, B. T. (2018). Tau induces blood vessel abnormalities and angiogenesis-related gene expression in P301L transgenic mice and human Alzheimer’s disease. Proceedings of the National Academy of Sciences, 115(6), E1289–E1298. doi:10.1073/pnas.1710329115

Benson, S., Pershadsingh, H. A., Ho, C. I., Chittiboyina, A., Desai, P., Pravenec, M., Qi, N., Wang, J., Avery, M. A., & Kurtz, T. W. (2004). Identification of Telmisartan as a Unique Angiotensin II Receptor Antagonist With Selective PPARgamma Modulating Activity. Hypertension, 43(5), 993–1002. doi:10.1161/01.HYP.0000123072.34629.57

Biffi, A., & Greenberg, S. M. (2011). Cerebral amyloid angiopathy: a systematic review. J Clin Neurol, 7(1), 1–9. 10.3988/jcn.2011.7.1.1

Bobotis, B. C., Braniff, O., Gargus, M., Akinluyi, E. T., Awogbindin, I. O., & Tremblay, M.-È. (2023). Sex differences of microglia in the healthy brain from embryonic development to adulthood and across lifestyle influences. Brain Research Bulletin, 202, 110752. 10.1016/j.brainresbull.2023.110752

Borzello, M., Ramirez, S., Treves, A., Lee, I., Scharfman, H., Stark, C., Knierim, J. J., & Rangel, L. M. (2023). Assessments of dentate gyrus function: discoveries and debates. Nature Reviews Neuroscience, 24(8), 502–517. 10.1038/s41583-023-00710-z

Boyle, P. A., Yu, L., Nag, S., Leurgans, S., Wilson, R. S., Bennett, D. A., & Schneider, J. A. (2015). Cerebral amyloid angiopathy and cognitive outcomes in community-based older persons. Neurology, 85(22), 1930–1936. 10.1212/wnl.0000000000002175

Cantero-Navarro, E., Fernández-Fernández, B., Ramos, A. M., Rayego-Mateos, S., Rodrigues-Diez, R. R., Sánchez-Niño, M. D., Sanz, A. B., Ruiz-Ortega, M., & Ortiz, A. (2021). Renin-angiotensin system and inflammation update. Molecular and cellular endocrinology, 529, 111254. 10.1016/j.mce.2021.111254

Casaletto, K. B., Nichols, E., Aslanyan, V., Simone, S. M., Rabin, J. S., La Joie, R., Brickman, A. M., Dams-O’Connor, K., Palta, P., Kumar, R. G., George, K. M., Satizabal, C. L., Schneider, J., & Pa, J. (2022). Sex-specific effects of microglial activation on Alzheimer’s disease proteinopathy in older adults. Brain, 145(10), 3536–3545. 10.1093/brain/awac257

Chen, X., Gao, R., Song, Y., Xu, T., Jin, L., Zhang, W., Chen, Z., Wang, H., Wu, W., Zhang, S., Zhang, G., Zhang, N., Chang, L., Liu, H., Li, H., & Wu, Y. (2023). Astrocytic AT1R deficiency ameliorates Aβ-induced cognitive deficits and synaptotoxicity through β-arrestin2 signaling. Progress in Neurobiology, 228, 102489. 10.1016/j.pneurobio.2023.102489

Chwalisz, B. K. (2021). Cerebral amyloid angiopathy and related inflammatory disorders. J Neurol Sci, 424, 117425. 10.1016/j.jns.2021.117425

Cooper, S. G., Souza, L. A. C., Worker, C. J., Gayban, A. J. B., Buller, S., Satou, R., & Feng Earley, Y. (2022). Renin-a in the Subfornical Organ Plays a Critical Role in the Maintenance of Salt-Sensitive Hypertension. Biomolecules, 12(9), 1169. https://www.mdpi.com/2218-273X/12/9/1169

Cosarderelioglu, C., Nidadavolu, L. S., George, C. J., Marx-Rattner, R., Powell, L., Xue, Q. L., Tian, J., Salib, J., Oh, E. S., Ferrucci, L., Dincer, P., Bennett, D. A., Walston, J. D., & Abadir, P. M. (2022). Higher Angiotensin II Type 1 Receptor Levels and Activity in the Postmortem Brains of Older Persons with Alzheimer’s Dementia. J Gerontol A Biol Sci Med Sci, 77(4), 664–672. 10.1093/gerona/glab376

Cosarderelioglu, C., Nidadavolu, L. S., George, C. J., Oh, E. S., Bennett, D. A., Walston, J. D., & Abadir, P. M. (2020). Brain Renin–Angiotensin System at the Intersect of Physical and Cognitive Frailty [Review]. Frontiers in Neuroscience, 14(981). 10.3389/fnins.2020.586314

Cozza, M., Amadori, L., & Boccardi, V. (2023). Exploring cerebral amyloid angiopathy: Insights into pathogenesis, diagnosis, and treatment. J Neurol Sci, 454, 120866. 10.1016/j.jns.2023.120866

Dahlof, B., Devereux, R. B., Kjeldsen, S. E., Julius, S., Beevers, G., de, F. U., Fyhrquist, F., Ibsen, H., Kristiansson, K., Lederballe-Pedersen, O., Lindholm, L. H., Nieminen, M. S., Omvik, P., Oparil, S., & Wedel, H. (2002). Cardiovascular morbidity and mortality in the Losartan Intervention For Endpoint reduction in hypertension study (LIFE): a randomised trial against atenolol. Lancet, 359(9311), 995–1003. PM:11937178 http://ac.els-cdn.com/S0140673602080893/1-s2.0-S0140673602080893-main.pdf?_tid=b5ce37e0-648d-11e3-9f34-00000aab0f27&acdnat=1387004735_8c93572d79fd1598dedfa18afc0fe789 (NOT IN FILE)

Daugherty, A., Rateri, D., Hong, L., & Balakrishnan, A. (2009). Measuring blood pressure in mice using volume pressure recording, a tail-cuff method. J Vis Exp(27). 10.3791/1291

Davis, J., Xu, F., Deane, R., Romanov, G., Previti, M. L., Zeigler, K., Zlokovic, B. V., & Van Nostrand, W. E. (2004). Early-onset and robust cerebral microvascular accumulation of amyloid beta-protein in transgenic mice expressing low levels of a vasculotropic Dutch/Iowa mutant form of amyloid beta-protein precursor. J Biol Chem, 279(19), 20296–20306. 10.1074/jbc.M312946200

de Gasparo, M., Catt, K. J., Inagami, T., Wright, J. W., & Unger, T. (2000). International union of pharmacology. XXIII. The angiotensin II receptors. Pharmacol.Rev., 52(3), 415–472. PM:10977869 http://pharmrev.aspetjournals.org/content/52/3/415.full.pdf (NOT IN FILE)

Delafontaine-Martel, P., Lefebvre, J., Damseh, R., Castonguay, A., Tardif, P., & Lesage, F. (2018). Large scale serial two-photon microscopy to investigate local vascular changes in whole rodent brain models of Alzheimer’s disease. Proc.SPIE,

Delage, C. I., Šimončičová, E., & Tremblay, M.-È. (2021). Microglial heterogeneity in aging and Alzheimer’s disease: Is sex relevant? J Pharmacol Sci, 146(3), 169–181. 10.1016/j.jphs.2021.03.006

DeSimone, C. V., Graff-Radford, J., El-Harasis, M. A., Rabinstein, A. A., Asirvatham, S. J., & Holmes, D. R., Jr. (2017). Cerebral Amyloid Angiopathy: Diagnosis, Clinical Implications, and Management Strategies in Atrial Fibrillation. J Am Coll Cardiol, 70(9), 1173–1182. 10.1016/j.jacc.2017.07.724

Filon, J. R., Intorcia, A. J., Sue, L. I., Vazquez Arreola, E., Wilson, J., Davis, K. J., Sabbagh, M. N., Belden, C. M., Caselli, R. J., Adler, C. H., Woodruff, B. K., Rapscak, S. Z., Ahern, G. L., Burke, A. D., Jacobson, S., Shill, H. A., Driver-Dunckley, E., Chen, K., Reiman, E. M., … Serrano, G. E. (2016). Gender Differences in Alzheimer Disease: Brain Atrophy, Histopathology Burden, and Cognition. Journal of Neuropathology & Experimental Neurology, 75(8), 748–754. 10.1093/jnen/nlw047

Fisher, R. A., Miners, J. S., & Love, S. (2022). Pathological changes within the cerebral vasculature in Alzheimer’s disease: New perspectives. Brain Pathol, 32(6), e13061. 10.1111/bpa.13061

Franklin, K. B. J., & Paxinos, G. (1997). *The Mouse Brain in Stereotaxic Coordinates*. Academic Press.

Gao, Y., Li, W., Liu, Y., Wang, Y., Zhang, J., Li, M., & Bu, M. (2018). Effect of Telmisartan on Preventing Learning and Memory Deficits Via Peroxisome Proliferator-Activated Receptor-γ in Vascular Dementia Spontaneously Hypertensive Rats. Journal of Stroke and Cerebrovascular Diseases, 27(2), 277–285. 10.1016/j.jstrokecerebrovasdis.2017.01.025

Gebre, A. K., Altaye, B. M., Atey, T. M., Tuem, K. B., & Berhe, D. F. (2018). Targeting Renin-Angiotensin System Against Alzheimer’s Disease. Front Pharmacol, 9, 440. 10.3389/fphar.2018.00440

Gireud-Goss, M., Mack, A. F., McCullough, L. D., & Urayama, A. (2021). Cerebral Amyloid Angiopathy and Blood-Brain Barrier Dysfunction. The Neuroscientist : a review journal bringing neurobiology, neurology and psychiatry, 27(6), 668–684. 10.1177/1073858420954811

Giuliani, A., Sivilia, S., Baldassarro, V. A., Gusciglio, M., Lorenzini, L., Sannia, M., Calzà, L., & Giardino, L. (2019). Age-Related Changes of the Neurovascular Unit in the Cerebral Cortex of Alzheimer Disease Mouse Models: A Neuroanatomical and Molecular Study. Journal of Neuropathology & Experimental Neurology, 78(2), 101–112. 10.1093/jnen/nly125

Gonzalez-Villalobos, R. A., Satou, R., Seth, D. M., Semprun-Prieto, L. C., Katsurada, A., Kobori, H., & Navar, L. G. (2009). Angiotensin-converting enzyme-derived angiotensin II formation during angiotensin II-induced hypertension. Hypertension, 53(2), 351–355. 10.1161/hypertensionaha.108.124511

Han, J., Fan, Y., Zhou, K., Blomgren, K., & Harris, R. A. (2021). Uncovering sex differences of rodent microglia. Journal of Neuroinflammation, 18(1), 74. 10.1186/s12974-021-02124-z

Haraguchi, T., Iwasaki, K., Takasaki, K., Uchida, K., Naito, T., Nogami, A., Kubota, K., Shindo, T., Uchida, N., Katsurabayashi, S., Mishima, K., Nishimura, R., & Fujiwara, M. (2010). Telmisartan, a partial agonist of peroxisome proliferator-activated receptor γ, improves impairment of spatial memory and hippocampal apoptosis in rats treated with repeated cerebral ischemia. Brain Research, 1353, 125–132. 10.1016/j.brainres.2010.07.017

Hu, Y. T., Chen, X. L., Zhang, Y. N., McGurran, H., Stormmesand, J., Breeuwsma, N., Sluiter, A., Zhao, J., Swaab, D., & Bao, A. M. (2023). Sex differences in hippocampal β-amyloid accumulation in the triple-transgenic mouse model of Alzheimer’s disease and the potential role of local estrogens. Front Neurosci, 17, 1117584. 10.3389/fnins.2023.1117584

Husain, A., Wilk, D., Smeby, R. R., Dzau, V. J., & Bumpus, F. M. (1984). Isorenin in dog brain and other tissues. Clin.Exp.Hypertens.A, 6(10-11), 1795–1799. PM:6398138 (NOT IN FILE)

Jankowsky, J. L., Melnikova, T., Fadale, D. J., Xu, G. M., Slunt, H. H., Gonzales, V., Younkin, L. H., Younkin, S. G., Borchelt, D. R., & Savonenko, A. V. (2005). Environmental enrichment mitigates cognitive deficits in a mouse model of Alzheimer’s disease. J Neurosci, 25(21), 5217–5224. 10.1523/jneurosci.5080-04.2005

Kara, F., van Dongen, E. S., Schliebs, R., van Buchem, M. A., de Groot, H. J. M., & Alia, A. (2012). Monitoring blood flow alterations in the Tg2576 mouse model of Alzheimer’s disease by in vivo magnetic resonance angiography at 17.6T. Neuroimage, 60(2), 958–966. 10.1016/j.neuroimage.2011.12.055

Kehoe, P. G., Miners, S., & Love, S. (2009). Angiotensins in Alzheimer’s disease - friend or foe? Trends Neurosci, 32(12), 619–628. 10.1016/j.tins.2009.07.006

Kehoe, P. G., & Wilcock, G. K. (2007). Is inhibition of the renin-angiotensin system a new treatment option for Alzheimer’s disease? Lancet Neurol., 6(4), 373–378. PM:17362841 (NOT IN FILE)

Kitanishi, T., Umaba, R., & Mizuseki, K. (2021). Robust information routing by dorsal subiculum neurons. Science Advances, 7(11), eabf1913. doi:10.1126/sciadv.abf1913

Kodama, L., & Gan, L. (2019). Do Microglial Sex Differences Contribute to Sex Differences in Neurodegenerative Diseases? Trends Mol Med, 25(9), 741–749. 10.1016/j.molmed.2019.05.001

Koran, M. E. I., Wagener, M., & Hohman, T. J. (2017). Sex differences in the association between AD biomarkers and cognitive decline. Brain Imaging Behav, 11(1), 205–213. 10.1007/s11682-016-9523-8

Kozberg, M. G., Perosa, V., Gurol, M. E., & van Veluw, S. J. (2021). A practical approach to the management of cerebral amyloid angiopathy. Int J Stroke, 16(4), 356–369. 10.1177/1747493020974464

Kuber, B., Fadnavis, M., & Chatterjee, B. (2023). Role of angiotensin receptor blockers in the context of Alzheimer’s disease. Fundam Clin Pharmacol, 37(3), 429–445. 10.1111/fcp.12872

Lee, H. W., Kim, S., Jo, Y., Kim, Y., Ye, B. S., & Yu, Y. M. (2023). Neuroprotective effect of angiotensin II receptor blockers on the risk of incident Alzheimer’s disease: A nationwide population-based cohort study. Frontiers in aging neuroscience, 15, 1137197. 10.3389/fnagi.2023.1137197

Lennon, M. J., Lipnicki, D. M., Lam, B. C. P., Crawford, J. D., Schutte, A. E., Peters, R., Rydberg-Sterner, T., Najar, J., Skoog, I., Riedel-Heller, S. G., Röhr, S., Pabst, A., Lobo, A., De-la-Cámara, C., Lobo, E., Lipton, R. B., Katz, M. J., Derby, C. A., Kim, K. W., … Sachdev, P. S. (2024). Blood Pressure, Antihypertensive Use, and Late-Life Alzheimer and Non-Alzheimer Dementia Risk: An Individual Participant Data Meta-Analysis. Neurology, 103(5), e209715. 10.1212/wnl.0000000000209715

Li, W., Zhang, J. W., Lu, F., Ma, M. M., Wang, J. Q., Suo, A. Q., Bai, Y. Y., & Liu, H. Q. (2012). [Effects of telmisartan on the level of Aβ1-42, interleukin-1β, tumor necrosis factor α and cognition in hypertensive patients with Alzheimer’s disease]. Zhonghua Yi Xue Za Zhi, 92(39), 2743–2746.

Loera-Valencia, R., Eroli, F., Garcia-Ptacek, S., & Maioli, S. (2021). Brain Renin–Angiotensin System as Novel and Potential Therapeutic Target for Alzheimer’s Disease. International Journal of Molecular Sciences, 22(18), 10139.

Lueptow, L. M. (2017). Novel Object Recognition Test for the Investigation of Learning and Memory in Mice. J Vis Exp(126). 10.3791/55718

Ma, A. P., Robertson, S. G., & Glass, B. D. (2022). Telmisartan Tablets Repackaged into Dose Administration Aids: Physicochemical Stability under Tropical Conditions. Pharmaceutics, 14(8). 10.3390/pharmaceutics14081667

Maniskas, M. E., Mack, A. F., Morales-Scheihing, D., Finger, C., Zhu, L., Paulter, R., Urayama, A., McCullough, L. D., & Manwani, B. (2021). Sex differences in a murine model of Cerebral Amyloid Angiopathy. Brain Behav Immun Health, 14, 100260. 10.1016/j.bbih.2021.100260

Mendel, T. A., Wierzba-Bobrowicz, T., Lewandowska, E., Stępień, T., & Szpak, G. M. (2013). The development of cerebral amyloid angiopathy in cerebral vessels. A review with illustrations based upon own investigated post mortem cases. Pol J Pathol, 64(4), 260–267. 10.5114/pjp.2013.39334

Merlo, S., Spampinato, S. F., Caruso, G. I., & Sortino, M. A. (2020). The Ambiguous Role of Microglia in Aβ Toxicity: Chances for Therapeutic Intervention. Curr Neuropharmacol, 18(5), 446–455. 10.2174/1570159x18666200131105418

Miao, J., Xu, F., Davis, J., Otte-Höller, I., Verbeek, M. M., & Van Nostrand, W. E. (2005). Cerebral microvascular amyloid beta protein deposition induces vascular degeneration and neuroinflammation in transgenic mice expressing human vasculotropic mutant amyloid beta precursor protein. Am J Pathol, 167(2), 505–515. 10.1016/s0002-9440(10)62993-8

Miners, J. S., Ashby, E., Van, H. Z., Chalmers, K. A., Palmer, L. E., Love, S., & Kehoe, P. G. (2008). Angiotensin-converting enzyme (ACE) levels and activity in Alzheimer’s disease, and relationship of perivascular ACE-1 to cerebral amyloid angiopathy. Neuropathol.Appl.Neurobiol., 34(2), 181–193. PM:17973905 http://onlinelibrary.wiley.com/store/10.1111/j.1365-2990.2007.00885.x/asset/j.1365-2990.2007.00885.x.pdf?v=1&t=hp4lhe7o&s=fb555ee5cb6b9e3b8ef1b970a135fb5ee275246d (NOT IN FILE)

Mogi, M., Li, J. M., Tsukuda, K., Iwanami, J., Min, L. J., Sakata, A., Fujita, T., Iwai, M., & Horiuchi, M. (2008). Telmisartan prevented cognitive decline partly due to PPAR-gamma activation. Biochem Biophys Res Commun, 375(3), 446–449. 10.1016/j.bbrc.2008.08.032

Naessens, D. M. P., de Vos, J., Richard, E., Wilhelmus, M. M. M., Jongenelen, C. A. M., Scholl, E. R., van der Wel, N. N., Heijst, J. A., Teunissen, C. E., Strijkers, G. J., Coolen, B. F., VanBavel, E., & Bakker, E. N. T. P. (2023). Effect of long-term antihypertensive treatment on cerebrovascular structure and function in hypertensive rats. Scientific Reports, 13(1), 3481. 10.1038/s41598-023-30515-0

Nebel, R. A., Aggarwal, N. T., Barnes, L. L., Gallagher, A., Goldstein, J. M., Kantarci, K., Mallampalli, M. P., Mormino, E. C., Scott, L., Yu, W. H., Maki, P. M., & Mielke, M. M. (2018). Understanding the impact of sex and gender in Alzheimer’s disease: A call to action. Alzheimers Dement, 14(9), 1171–1183. 10.1016/j.jalz.2018.04.008

Nikolajsen, G. N., Kotynski, K. A., Jensen, M. S., & West, M. J. (2015). Quantitative analysis of the capillary network of aged APPswe/PS1dE9 transgenic mice. Neurobiology Of Aging, 36(11), 2954–2962. 10.1016/j.neurobiolaging.2015.08.004

O’Mara, S. (2005). The subiculum: what it does, what it might do, and what neuroanatomy has yet to tell us. J Anat, 207(3), 271–282. 10.1111/j.1469-7580.2005.00446.x

Ouk, M., Wu, C. Y., Rabin, J. S., Jackson, A., Edwards, J. D., Ramirez, J., Masellis, M., Swartz, R. H., Herrmann, N., Lanctôt, K. L., Black, S. E., & Swardfager, W. (2021). The use of angiotensin-converting enzyme inhibitors vs. angiotensin receptor blockers and cognitive decline in Alzheimer’s disease: the importance of blood-brain barrier penetration and APOE ε4 carrier status. Alzheimers Res Ther, 13(1), 43. 10.1186/s13195-021-00778-8

Oveisgharan, S., Arvanitakis, Z., Yu, L., Farfel, J., Schneider, J. A., & Bennett, D. A. (2018). Sex differences in Alzheimer’s disease and common neuropathologies of aging. Acta Neuropathol, 136(6), 887–900. 10.1007/s00401-018-1920-1

Pang, T., Sun, L. X., Wang, T., Jiang, Z. Z., Liao, H., & Zhang, L. Y. (2014). Telmisartan protects central neurons against nutrient deprivation-induced apoptosis in vitro through activation of PPARgamma and the Akt/GSK-3beta pathway. Acta Pharmacol Sin, 35(6), 727–737. 10.1038/aps.2013.199

Paul, J. R., Munir, H. A., van Groen, T., & Gamble, K. L. (2018). Behavioral and SCN neurophysiological disruption in the Tg-SwDI mouse model of Alzheimer’s disease. Neurobiol Dis, 114, 194–200. 10.1016/j.nbd.2018.03.007

Rea, F., Corrao, G., & Mancia, G. (2024). Risk of Dementia During Antihypertensive Drug Therapy in the Elderly. J Am Coll Cardiol, 83(13), 1194–1203. 10.1016/j.jacc.2024.01.030

Restrepo, Y. M., Noto, N. M., & Speth, R. C. (2022). CGP42112: the full AT2 receptor agonist and its role in the renin-angiotensin-aldosterone system: no longer misunderstood. Clin Sci (Lond), 136(21), 1513–1533. 10.1042/cs20220261

Ribeiro, V. T., de Souza, L. C., & Simões, E. S. A. C. (2020). Renin-Angiotensin System and Alzheimer’s Disease Pathophysiology: From the Potential Interactions to Therapeutic Perspectives. Protein Pept Lett, 27(6), 484–511. 10.2174/0929866527666191230103739

Robison, L. S., Francis, N., Popescu, D. L., Anderson, M. E., Hatfield, J., Xu, F., Anderson, B. J., Van Nostrand, W. E., & Robinson, J. K. (2020). Environmental Enrichment: Disentangling the Influence of Novelty, Social, and Physical Activity on Cerebral Amyloid Angiopathy in a Transgenic Mouse Model. Int J Mol Sci, 21(3). 10.3390/ijms21030843

Robison, L. S., Popescu, D. L., Anderson, M. E., Francis, N., Hatfield, J., Sullivan, J. K., Beigelman, S. I., Xu, F., Anderson, B. J., Van Nostrand, W. E., & Robinson, J. K. (2019). Long-term voluntary wheel running does not alter vascular amyloid burden but reduces neuroinflammation in the Tg-SwDI mouse model of cerebral amyloid angiopathy. J Neuroinflammation, 16(1), 144. 10.1186/s12974-019-1534-0

Rodriguez-Lopez, A., Torres-Paniagua, A. M., Acero, G., Díaz, G., & Gevorkian, G. (2023). Increased TSPO expression, pyroglutamate-modified amyloid beta (AβN3(pE)) accumulation and transient clustering of microglia in the thalamus of Tg-SwDI mice. Journal of Neuroimmunology, 382, 578150. 10.1016/j.jneuroim.2023.578150

Rosas-Hernandez, H., Cuevas, E., Raymick, J. B., Robinson, B. L., & Sarkar, S. (2020). Impaired Amyloid Beta Clearance and Brain Microvascular Dysfunction are Present in the Tg-SwDI Mouse Model of Alzheimer’s Disease. Neuroscience, 440, 48–55. 10.1016/j.neuroscience.2020.05.024

Saavedra, J. M. (2021). Angiotensin Receptor Blockers Are Not Just for Hypertension Anymore. Physiology (Bethesda), 36(3), 160–173. 10.1152/physiol.00036.2020

Saito, S., Tanaka, M., Satoh-Asahara, N., Carare, R. O., & Ihara, M. (2021). Taxifolin: A Potential Therapeutic Agent for Cerebral Amyloid Angiopathy. Front Pharmacol, 12, 643357. 10.3389/fphar.2021.643357

Santisteban, M. M., Iadecola, C., & Carnevale, D. (2023). Hypertension, Neurovascular Dysfunction, and Cognitive Impairment. Hypertension, 80(1), 22–34. 10.1161/hypertensionaha.122.18085

Santos, R. A. S., Sampaio, W. O., Alzamora, A. C., Motta-Santos, D., Alenina, N., Bader, M., & Campagnole-Santos, M. J. (2018). The ACE2/Angiotensin-(1-7)/MAS Axis of the Renin-Angiotensin System: Focus on Angiotensin-(1-7). Physiol Rev, 98(1), 505–553. 10.1152/physrev.00023.2016

Savaskan, E., Hock, C., Olivieri, G., Bruttel, S., Rosenberg, C., Hulette, C., & Muller-Spahn, F. (2001). Cortical alterations of angiotensin converting enzyme, angiotensin II and AT1 receptor in Alzheimer’s dementia. Neurobiology Of Aging, 22(4), 541–546. PM:11445253 http://ac.els-cdn.com/S0197458000002591/1-s2.0-S0197458000002591-main.pdf?_tid=3e814eae-6387-11e3-b1b5-00000aacb35d&acdnat=1386892007_586758347d5501b53da27f37c87f751e (NOT IN FILE)

Sessle, B. J. (2016). Chapter 1 - The Biological Basis of a Functional Occlusion: The Neural Framework. In I. Klineberg & S. E. Eckert (Eds.), Functional Occlusion in Restorative Dentistry and Prosthodontics (pp. 3–22). Mosby. 10.1016/B978-0-7234-3809-0.00001-2

Shindo, T., Takasaki, K., Uchida, K., Onimura, R., Kubota, K., Uchida, N., Irie, K., Katsurabayashi, S., Mishima, K., Nishimura, R., Fujiwara, M., & Iwasaki, K. (2012). Ameliorative effects of telmisartan on the inflammatory response and impaired spatial memory in a rat model of Alzheimer’s disease incorporating additional cerebrovascular disease factors. Biol Pharm Bull, 35(12), 2141–2147. 10.1248/bpb.b12-00387

Sin, M. K., Zamrini, E., Ahmed, A., Nho, K., & Hajjar, I. (2023). Anti-Amyloid Therapy, AD, and ARIA: Untangling the Role of CAA. J Clin Med, 12(21). 10.3390/jcm12216792

Singh, B., Sharma, B., Jaggi, A. S., & Singh, N. (2013). Attenuating effect of lisinopril and telmisartan in intracerebroventricular streptozotocin induced experimental dementia of Alzheimer’s disease type: possible involvement of PPAR-gamma agonistic property. J Renin Angiotensin Aldosterone Syst, 14(2), 124–136. 10.1177/1470320312459977 10.1177/1470320312459977. Epub 2012 Oct 11.

Siracusa, R., Fusco, R., & Cuzzocrea, S. (2019). Astrocytes: Role and Functions in Brain Pathologies. Front Pharmacol, 10, 1114. 10.3389/fphar.2019.01114

Speth, R. C. (2022). Renin-Angiotensin-Aldosterone System. In T. Kenakin (Ed.), Comprehensive Pharmacology. (Vol. 4, pp. 528–569). Elsevier. 10.1016/B978-0-12-820472-6.00160-2.

Stornetta, R. L., Hawelu-Johnson, C. L., Guyenet, P. G., & Lynch, K. R. (1988). Astrocytes synthesize angiotensinogen in brain. Science, 242, 1444–1446. http://media.proquest.com/media/pq/classic/doc/1789381/fmt/pi/rep/NONE?hl=&cit%3Aauth=Stornetta%2C+Ruth+L%3BHawelu-Johnson%2C+Charlyn+L%3BGuyenet%2C+Patrice+G%3BLynch%2C+Kevin+R&cit% 3Atitle=Astrocytes+Synthesize+Angiotensinogen+in+Brain&cit%3Apub=Science&cit% 3Avol=242&cit%3Aiss=4884&cit%3Apg=1444&cit%3Adate=Dec+9%2C+1988&ic=tru e&cit%3Aprod=ProQuest&_a=ChgyMDEzMTIxNDIwMTU0MjczMTo3NjA2MDUSBj ExNTk2NBoKT05FX1NFQVJDSCIMMTM3LjUyLjM0Ljk2KgQxMjU2MgkyMTM1M zM1ODg6DURvY3VtZW50SW1hZ2VCATBSBk9ubGluZVoCRlRiA1BGVGoKMTk4 OC8xMi8wOXIKMTk4OC8xMi8wOXoAggEyUC0xMDA3MDY3LTQwNzY1LUNVU 1RPTUVSLTEwMDAwMDM5LzEwMDAwMTU1LTEwNjUwOTGSAQZPbmxpbmXKAQdFbmROb3Rl0gESU2Nob2xhcmx5IEpvdXJuYWxzmgIHUHJlUGFpZKoCKE9TOk VNUy1QZGZEb2NWaWV3QmFzZS1nZXRNZWRpYVVybEZvckl0ZW2yAgC6AgDKAg9BcnRpY2xlfEZlYXR1cmU%3D&_s=Dz6HFWDPtEo35zdoWkF%2B8teBdD0%3D (IN FILE)

Strittmatter, S. M., Thiele, E. A., Kapiloff, M. S., & Snyder, S. H. (1985). A rat brain isozyme of angiotensin-converting enzyme: unique specificity for amidated peptide substrates. J.Biol.Chem., 260, 9825–9832. http://www.jbc.org/content/260/17/9825.full.pdf (IN FILE)

Sun, D., Mei, L., & Xiong, W.-C. (2023). Dorsal Dentate Gyrus, a Key Regulator for Mood and Psychiatric Disorders. Biological Psychiatry, 93(12), 1071–1080. 10.1016/j.biopsych.2023.01.005

Sveikata, L., Charidimou, A., & Viswanathan, A. (2022). Vessels Sing Their ARIAs: The Role of Vascular Amyloid in the Age of Aducanumab. Stroke, 53(1), 298–302. 10.1161/strokeaha.121.036873

Taylor, X., Cisternas, P., You, Y., You, Y., Xiang, S., Marambio, Y., Zhang, J., Vidal, R., & Lasagna-Reeves, C. A. (2020). A1 reactive astrocytes and a loss of TREM2 are associated with an early stage of pathology in a mouse model of cerebral amyloid angiopathy. J Neuroinflammation, 17(1), 223. 10.1186/s12974-020-01900-7

Tchekalarova, J., Ivanova, P., & Krushovlieva, D. (2024). Age-Related Effects of AT1 Receptor Antagonist Losartan on Cognitive Decline in Spontaneously Hypertensive Rats. Int J Mol Sci, 25(13). 10.3390/ijms25137340

Thal, D. R., Capetillo-Zarate, E., Larionov, S., Staufenbiel, M., Zurbruegg, S., & Beckmann, N. (2009). Capillary cerebral amyloid angiopathy is associated with vessel occlusion and cerebral blood flow disturbances. Neurobiology Of Aging, 30(12), 1936–1948. 10.1016/j.neurobiolaging.2008.01.017

Torrico, T. J., & Munakomi, S. (2025). Neuroanatomy, Thalamus. In StatPearls. StatPearls Publishing Copyright © 2025, StatPearls Publishing LLC.

Tsukuda, K., Mogi, M., Iwanami, J., Min, L. J., Sakata, A., Jing, F., Iwai, M., & Horiuchi, M. (2009). Cognitive deficit in amyloid-beta-injected mice was improved by pretreatment with a low dose of telmisartan partly because of peroxisome proliferator-activated receptor-gamma activation. Hypertension, 54(4), 782–787. 10.1161/hypertensionaha.109.136879

Vila-Castelar, C., Akinci, M., Palpatzis, E., Aguilar-Dominguez, P., Operto, G., Kollmorgen, G., Quijano-Rubio, C., Blennow, K., Zetterberg, H., Falcon, C., Fauria, K., Gispert, J. D., Grau-Rivera, O., Suárez-Calvet, M., Arenaza-Urquijo, E. M., Anastasi, F., Beteta, A., Brugulat-Serrat, A., Cacciaglia, R., … for the, A. s. (2024). Sex/gender effects of glial reactivity on preclinical Alzheimer’s disease pathology. Molecular Psychiatry. 10.1038/s41380-024-02753-9

Villapol, S., Janatpour, Z. C., Affram, K. O., & Symes, A. J. (2023). The Renin Angiotensin System as a Therapeutic Target in Traumatic Brain Injury. Neurotherapeutics, 20(6), 1565–1591. 10.1007/s13311-023-01435-8

Wang, Y., Thatcher, S. E., & Cassis, L. A. (2017). Measuring Blood Pressure Using a Noninvasive Tail Cuff Method in Mice. Methods in Molecular Biology, 1614, 69–73. 10.1007/978-1-4939-7030-8_6

Washida, K., Ihara, M., Nishio, K., Fujita, Y., Maki, T., Yamada, M., Takahashi, J., Wu, X., Kihara, T., Ito, H., Tomimoto, H., & Takahashi, R. (2010). Nonhypotensive dose of telmisartan attenuates cognitive impairment partially due to peroxisome proliferator-activated receptor-gamma activation in mice with chronic cerebral hypoperfusion. Stroke, 41(8), 1798–1806. 10.1161/strokeaha.110.583948

Weller, R. O., Subash, M., Preston, S. D., Mazanti, I., & Carare, R. O. (2008). Perivascular drainage of amyloid-beta peptides from the brain and its failure in cerebral amyloid angiopathy and Alzheimer’s disease. Brain Pathol, 18(2), 253–266. 10.1111/j.1750-3639.2008.00133.x

Whiteford, J. R., De Rossi, G., & Woodfin, A. (2016). Mutually Supportive Mechanisms of Inflammation and Vascular Remodeling. Int Rev Cell Mol Biol, 326, 201–278. 10.1016/bs.ircmb.2016.05.001

Xu, F., Grande, A. M., Robinson, J. K., Previti, M. L., Vasek, M., Davis, J., & Van Nostrand, W. E. (2007). Early-onset subicular microvascular amyloid and neuroinflammation correlate with behavioral deficits in vasculotropic mutant amyloid beta-protein precursor transgenic mice. Neuroscience, 146(1), 98–107. 10.1016/j.neuroscience.2007.01.043

Xu, W., Xu, F., Anderson, M. E., Kotarba, A. E., Davis, J., Robinson, J. K., & Van Nostrand, W. E. (2014). Cerebral microvascular rather than parenchymal amyloid-β protein pathology promotes early cognitive impairment in transgenic mice. J Alzheimers Dis, 38(3), 621–632. 10.3233/jad-130758

Xu, X., Meng, T., Wen, Q., Tao, M., Wang, P., Zhong, K., & Shen, Y. (2020). Dynamic changes in vascular size and density in transgenic mice with Alzheimer’s disease. Aging (Albany NY), 12(17), 17224–17234. 10.18632/aging.103672

Xu, Y., Gao, W., Sun, Y., & Wu, M. (2023). New insight on microglia activation in neurodegenerative diseases and therapeutics. Front Neurosci, 17, 1308345. 10.3389/fnins.2023.1308345

Yasar, S., Xia, J., Yao, W., Furberg, C. D., Xue, Q. L., Mercado, C. I., Fitzpatrick, A. L., Fried, L. P., Kawas, C. H., Sink, K. M., Williamson, J. D., DeKosky, S. T., & Carlson, M. C. (2013). Antihypertensive drugs decrease risk of Alzheimer disease: Ginkgo Evaluation of Memory Study [WNL.0b013e3182a35228 pii; 10.1212/WNL.0b013e3182a35228 doi]. Neurology, 81(10), 896-903. PM:23911756 http://graphics.tx.ovid.com/ovftpdfs/FPDDNCOBIDHKOC00/fs047/ovft/live/gv024/00006114/00006114-201309030-00011.pdf (NOT IN FILE)

Zlokovic, B. V. (2011). Neurovascular pathways to neurodegeneration in Alzheimer’s disease and other disorders. Nature Reviews Neuroscience, 12(12), 723–738. 10.1038/nrn3114

