## Supplementary material for "Telmisartan and Lisinopril Show Potential Benefits in Rescuing Cognitive-Behavioral Function Despite Limited Improvements in Neuropathological Outcomes in Tg-SwDI Mice": Graphical Abstract

### 1 Animals

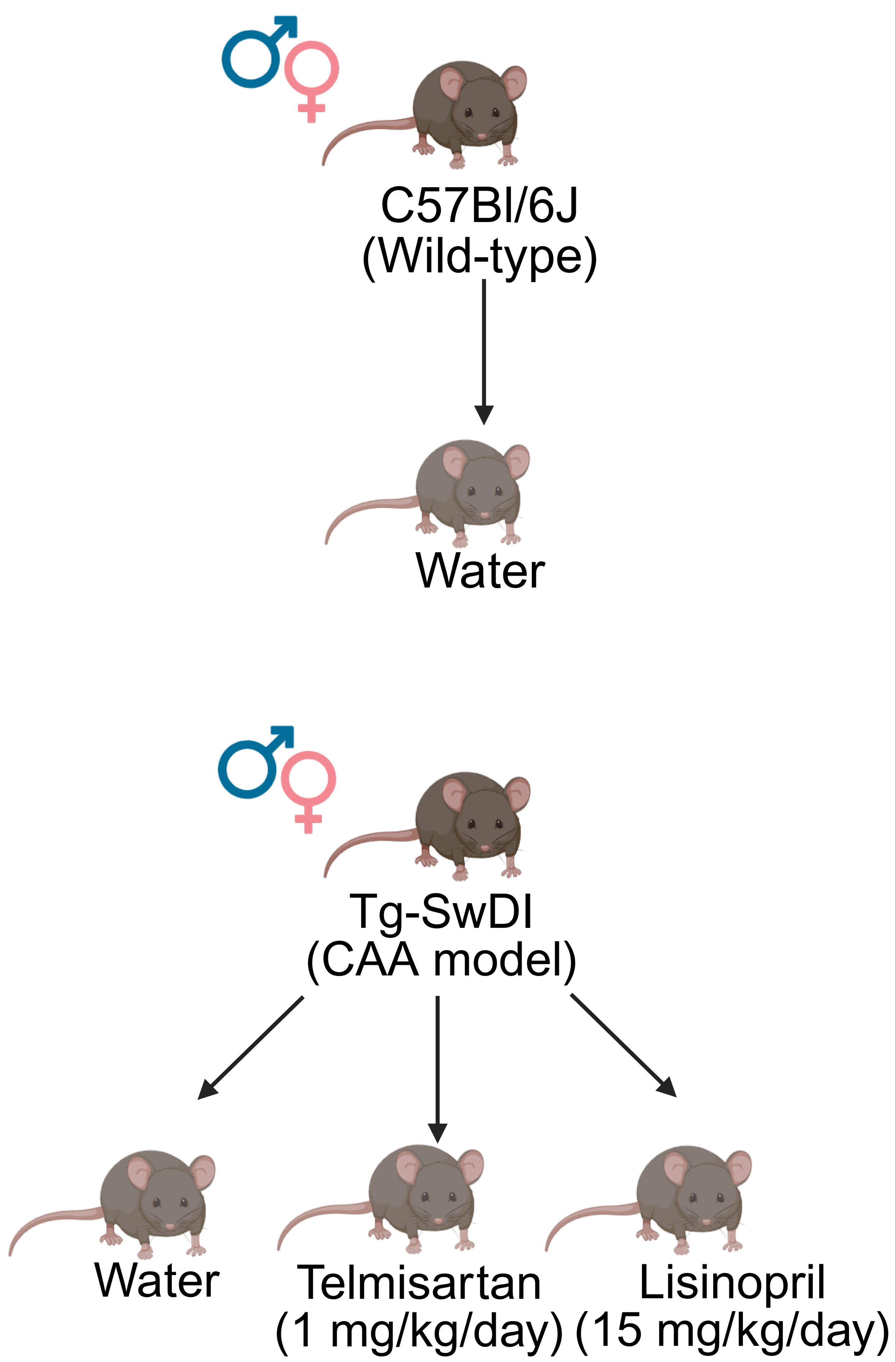


### 2 Behavior Testing

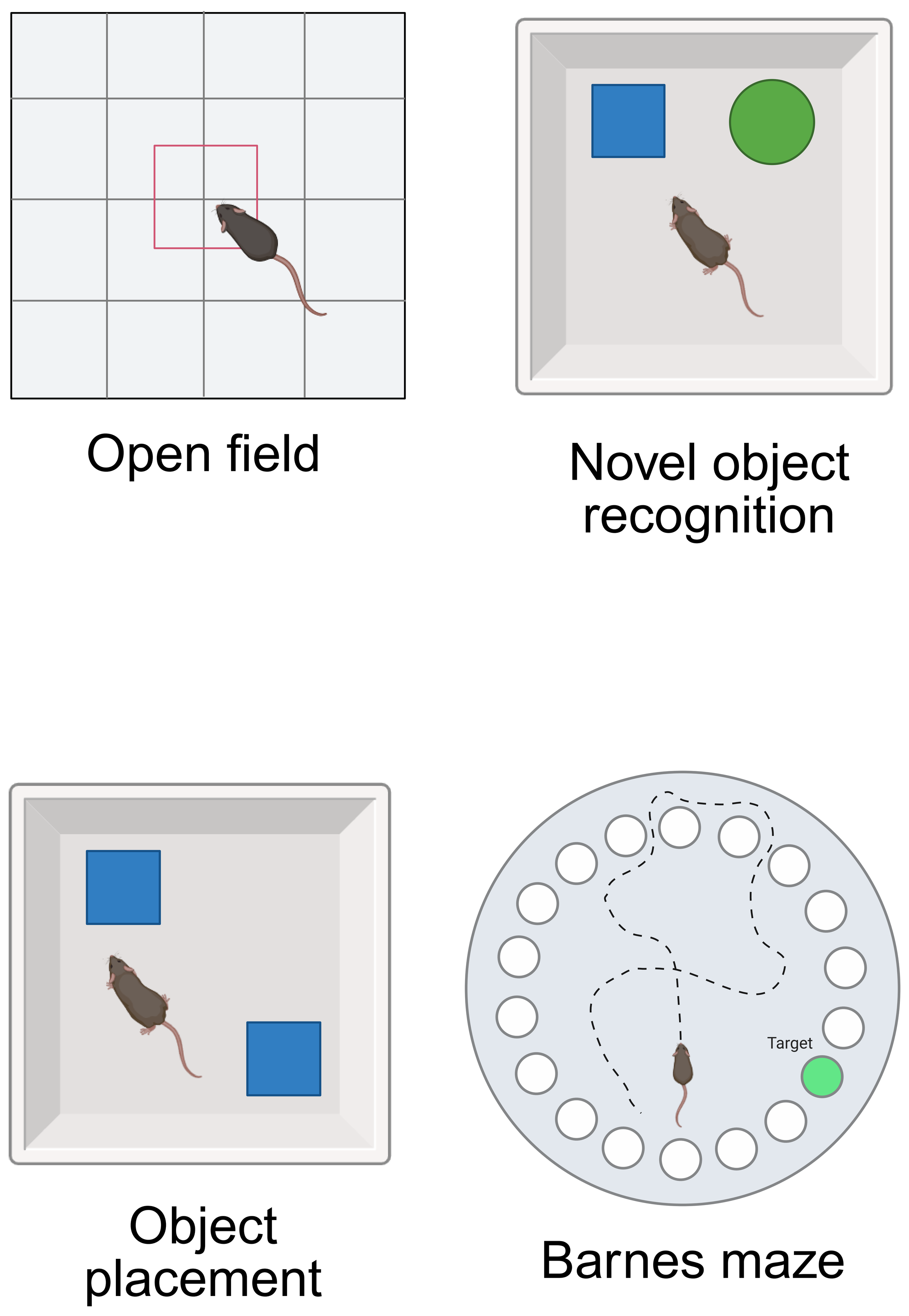


### 3 Histochemistry

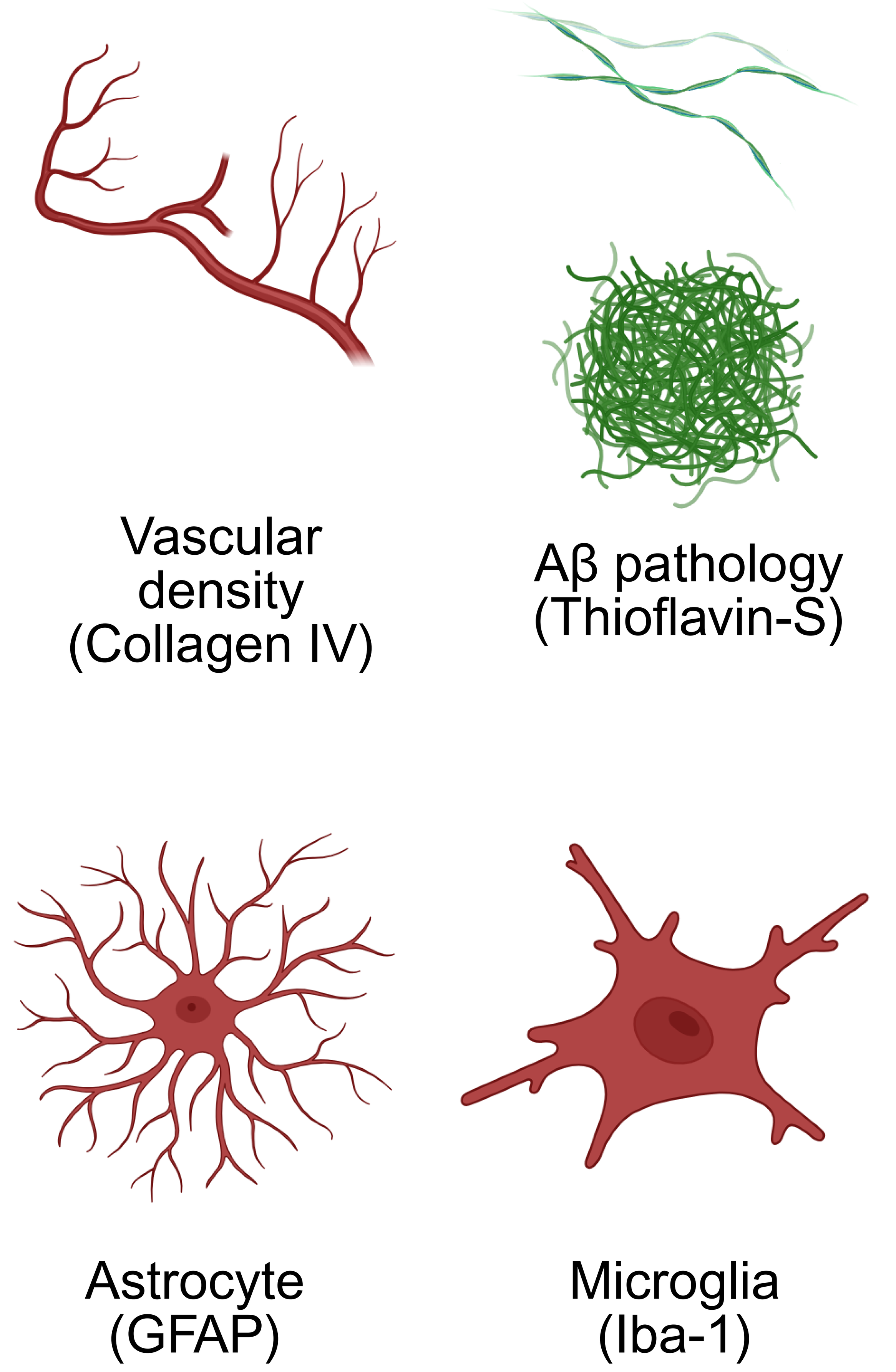


### 4 Findings

**TELMISARTAN**

- Partially improved object recognition and short- and long-term spatial memory
- Partially ameliorated vascular deficit in the ventrolateral thalamus (females), astrogliosis in the dentate gyrus (males), and microgliosis in the dorsal subiculum (females)

**LISINOPRIL**

- Improved short- and long-term spatial memory
- Partially ameliorated vascular deficit in the dentate gyrus and microgliosis in the sensorimotor cortex (females)
- Modestly increased Aβ pathology in the ventrolateral thalamus (males)

### Conclusion:

Telmisartan and lisinopril were able to rescue some cognitive-behavioral functions in early-stage-disease Tg-SwDI mice. However, no reductions in Aβ levels were observed, and limited improvements in vascular density and neuroinflammatory markers were detected. Moreover, some of the effects observed were sex-, behavioral task-, and brain region-specific.

*Created in BioRender. Speth, R. (2025) https://BioRender.com/21ku05a*
