## Supplementary material for "Telmisartan and Lisinopril Show Potential Benefits in Rescuing Cognitive-Behavioral Function Despite Limited Improvements in Neuropathological Outcomes in Tg-SwDI Mice": Figure 1 - Publication License

49 Spadina Ave. Suite 200
Toronto ON M5V 2J1 Canada
www.biorender.com

### Confirmation of Publication and Licensing Rights - Open Access

**April 23rd, 2025**

| | |
|---|---|
| ***Subscription Type:*** | *Individual - Academic* |
| ***Agreement number:*** | *XJ286LLWP1* |
| ***Publisher Name:*** | *Biorxiv* |
| ***Figure Title:*** | *Simplified representation of the activity of the vasculature and the brain RAS* |
| ***Citation to Use:*** | *Created in BioRender. Speth, R. (2025) https://BioRender.com/rg5y89w* |

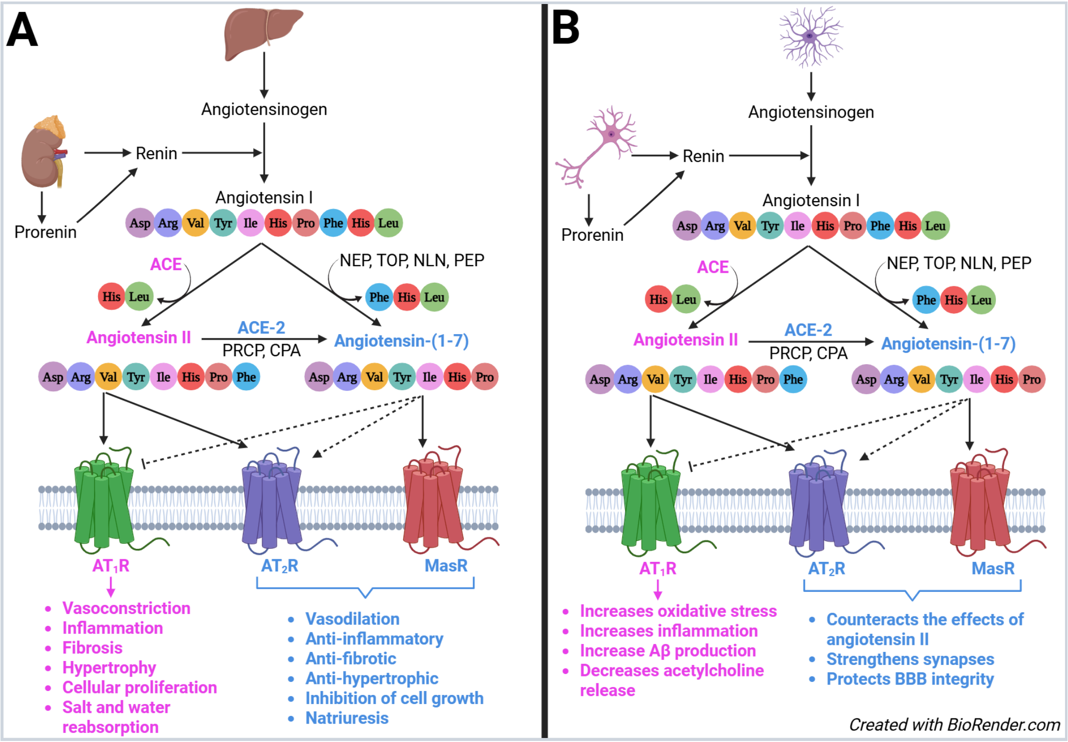


*For any questions regarding this document, or other questions about publishing with BioRender, please refer to our [BioRender Publication Guide](#), or contact BioRender Support at [](#).*
