## Supplementary material for "Telmisartan and Lisinopril Show Potential Benefits in Rescuing Cognitive-Behavioral Function Despite Limited Improvements in Neuropathological Outcomes in Tg-SwDI Mice": Glossary

*Alzheimer's disease (AD):* As defined by the National Institute on Aging (NIA), AD is the most common type of dementia, characterized by slow destruction of memory and thinking skills, as well as behavioral changes and ability to carry out simple tasks.

*Amyloid-beta (Aβ):* An oligomeric peptide that ranges from 39-43 amino acids in length, that arises from cleavage of the amyloid precursor protein.

*Amyloid-related imaging abnormality:* Abnormal differences observed in MRIs of brains, a morbidity often seen in patients taking anti-amyloid antibody drugs.

*Angiotensin-converting enzyme (ACE) inhibitor:* A class of drugs, commonly prescribed to treat hypertension, that inhibit the conversion of angiotensin I into angiotensin II.

*Angiotensin receptor blocker (ARB):* A class of drugs, commonly prescribed to treat hypertension, that block the AT_1_R, preventing angiotensin II from binding.

*Bregma:* The point where the coronal and sagittal sutures of the skull join.

*C57BL/6J:* The mouse strain used as the genetic background for the development of Tg-SwDI mice.

*Cerebral amyloid angiopathy (CAA):* A cerebrovascular disease characterized by the accumulation of Aβ.

*Renin-angiotensin system (RAS):* Hormone system responsible for regulating blood pressure, electrolyte balance, and fluid balance.

*Tail-cuff method:* A technique used to measure blood pressure in rodents, where a cuff is placed on the animal’s tail to measure changes in blood volume as the system inflates and deflates.

*Tg-SwDI:* A transgenic mouse model of CAA that carries the Dutch (E693Q), Iowa (D694N), and Swedish (K670N/M671L) mutations in the amyloid precursor protein.
